# Deep contrastive learning enables genome-wide virtual screening

**DOI:** 10.1101/2024.09.02.610777

**Authors:** Yinjun Jia, Bowen Gao, Jiaxin Tan, Jiqing Zheng, Xin Hong, Wenyu Zhu, Haichuan Tan, Yuan Xiao, Liping Tan, Hongyi Cai, Yanwen Huang, Zhiheng Deng, Xiangwei Wu, Yue Jin, Yafei Yuan, Jiekang Tian, Wei He, Weiying Ma, Yaqin Zhang, Wei Zhang, Lei Liu, Chuangye Yan, Yanyan Lan

## Abstract

Numerous protein-coding genes are associated with human diseases, yet approximately 90% of them lack targeted therapeutic intervention. While conventional computational methods, such as molecular docking, have facilitated the discovery of potential hit compounds, the development of genome-wide virtual screening against the expansive chemical space remains a formidable challenge. Here we introduce DrugCLIP, a novel framework that combines contrastive learning and dense retrieval to achieve rapid and accurate virtual screening. Compared to traditional docking methods, DrugCLIP improves the speed of virtual screening by up to seven orders of magnitude. In terms of performance, DrugCLIP not only surpasses docking and other deep learning-based methods across two standard benchmark datasets, but also demonstrates high efficacy in wet-lab experiments. Specifically, DrugCLIP successfully identified agonists with < 100 nM affinities for 5HT_2A_R, a key target in psychiatric diseases. For another target NET, whose structure is newly solved and not included in the training set, our method achieved a hit rate of 15%, with 12 diverse molecules exhibiting affinities better than bupropion. Additionally, two chemically novel inhibitors were validated by structure determination with Cryo-EM. Finally, a novel potential drug target TRIP12, with no experimental structures and inhibitors for reference, was used to challenge DrugCLIP. DrugCLIP achieved a hit rate of 17.5% by screening a pocket identified on an AlphaFold2-predicted structure, verified with multi-cycle SPR assays. Molecules with the highest affinities also showed a dose-dependent inhibition to the enzymatic function of TRIP12. Building on this foundation, we present the results of a pioneering trillion-scale genome-wide virtual screening, encompassing approximately 10,000 AlphaFold2 predicted proteins within the human genome and 500 million molecules from the ZINC and Enamine REAL database. This work provides an innovative perspective on drug discovery in the post-AlphaFold era, where comprehensive targeting of all disease-related proteins is within reach.

## Introduction

The human genome comprises approximately 20,000 protein-coding genes (*1*), many of which are related to a variety of diseases. Despite this, only about 10% of these genes have been successfully targeted by FDA-approved drugs or have documented small-molecule binders in the literature (*2*). This leaves a substantial portion of the druggable genome largely unexplored, representing a promising opportunity for therapeutic innovation. The scientific community is eager to translate biologically relevant targets into pharmaceutical breakthroughs. However, most researchers lack access to advanced high-throughput screening equipment or sufficient computational power to perform comprehensive virtual screenings. Additionally, proteins often function as parts of families or pathways, indicating that targeting single proteins may not always be the most effective strategy (*3, 4*). These limitations can significantly reduce the success rate of drug discovery, especially for new targets. Therefore, developing a comprehensive chemical database containing genome-wide virtual screening results would be an invaluable asset for the biomedical research community, with the potential to significantly accelerate the discovery of new drugs.

Given the impracticality of experimentally screening all human proteins, virtual screening has emerged as the only viable approach to tackle the vast number of potential targets. In classical computer-aided drug discovery (CADD), molecular docking serves as a foundational technique for target-based virtual screening. Despite advancements in simplified scoring functions, optimized algorithms, and hardware acceleration (*5–9*), molecular docking remains time-intensive, often requiring several seconds to minutes to evaluate each protein-ligand pair. For example, a recent large-scale docking campaign took two weeks to screen 1 billion molecules against a single target, even with the use of 10,000 CPU cores (*10*). As a result, the computational demands for genome-wide virtual screening are prohibitively high, rendering such efforts impractical with existing technologies.

Artificial intelligence holds great promise for drug discovery. Various deep learning methods have been developed for virtual screening, focusing on predicting ligand-receptor affinities (*11–13*). Yet, applying these methods to large-scale virtual screening still faces significant challenges. A primary issue is the inconsistency of affinity values due to heterogeneous experimental conditions (*14, 15*), which may negatively impact the performance of the trained model. Moreover, a notable distribution shift between training datasets and real-world testing scenarios hinders the generalizability of AI models, as real-world virtual screenings often involve a larger proportion of inactive molecules than those represented in the curated training sets (*16*). Additionally, the computational demands of deep learning models, with millions of parameters, pose a crucial bottleneck in inference speed, especially as chemical libraries and target numbers grow. Consequently, there is an urgent need for the development of more efficient and robust AI methodologies to effectively address these challenges.

In this work, we introduce DrugCLIP, a novel contrastive learning approach for virtual screening. Contrastive learning has demonstrated significant success in various applications like image-text retrieval (*17*), enzyme function annotation (*18*), and protein homology detection (*19*). The core innovation of DrugCLIP lies in its ability to distinguish potent binders from non-binding molecules with a given protein pocket by aligning their representations. This approach effectively mitigates the impact of noisy affinity labels and chemical library imbalances that have traditionally challenged virtual screening efforts. Moreover, the inference of DrugCLIP is highly efficient, achieving a speed improvement in several orders of magnitude.

Comprehensive *in silico* and wet-lab evaluations were conducted to assess the accuracy of the DrugCLIP model. Our model achieved state-of-the-art performance on two widely recognized virtual screening benchmarks, DUD-E (*20*) and LIT-PCBA (*21*), outperforming traditional docking-based screening methods and other deep neural networks. To further validate its performance, DrugCLIP was applied to screen molecules for three real-world targets: 5HT_2A_R (5-hydroxytryptamine receptor 2A), NET (norepinephrine transporter), and TRIP12 (Thyroid Hormone Receptor Interactor 12), while the last target, TRIP12, lacks experimental structures and inhibitors for reference. Remarkably, our model identified chemically diverse binders with adequate affinities, which were further validated through functional assays and structure determination. These results provide compelling evidence of the efficacy of our virtual screening method.

Finally, a genome-wide virtual screening was conducted using DrugCLIP on all human proteins predicted by AlphaFold2 (*22, 23*). In this process, we first define pockets for AlphaFold predictions with structure alignment (*24*), pocket detection software (*25*), and generative AI models. Next, we screened over 500 million drug-like molecules from the ZINC (*26, 27*) and Enamine REAL (*28*) databases against identified pockets. Notably, this unprecedented large-scale virtual screening was completed in just 24 hours on a single computing node equipped with 8 A100 GPUs. Lastly, we applied a CADD cluster-docking pipeline to select chemically diverse and physically proper molecules for each pocket. These result in a dataset containing over 2 million potential hits targeting more than 20,000 pockets from around 10,000 human proteins. To the best of our knowledge, this is the first virtual screening campaign to perform more than 10 trillion scoring operations on protein-ligand pairs, covering nearly half of the human genome. All molecules, scores, and poses have been made freely accessible at https://drug-the-whole-genome.yanyanlan.com, facilitating further research in drug discovery on a genome-wide scale.

## Results

### The design of the DrugCLIP model

Unlike previous machine learning models that relied on regression to directly predict protein-ligand affinity values, DrugCLIP (Fig. 1) redefines virtual screening as a dense retrieval task. The key innovation lies in its training objective, which aims to learn an aligned embedding space for protein pockets and molecules, encoded by separate neural networks. Vector similarity metrics can then be employed to reflect their binding probability. Using contrastive loss during training, the similarity between protein pockets and their binders (positive protein-ligand pairs) is maximized, whereas the similarity between protein pockets and molecules binding to other targets (negative protein-ligand pairs) is minimized.

**Fig. 1.**
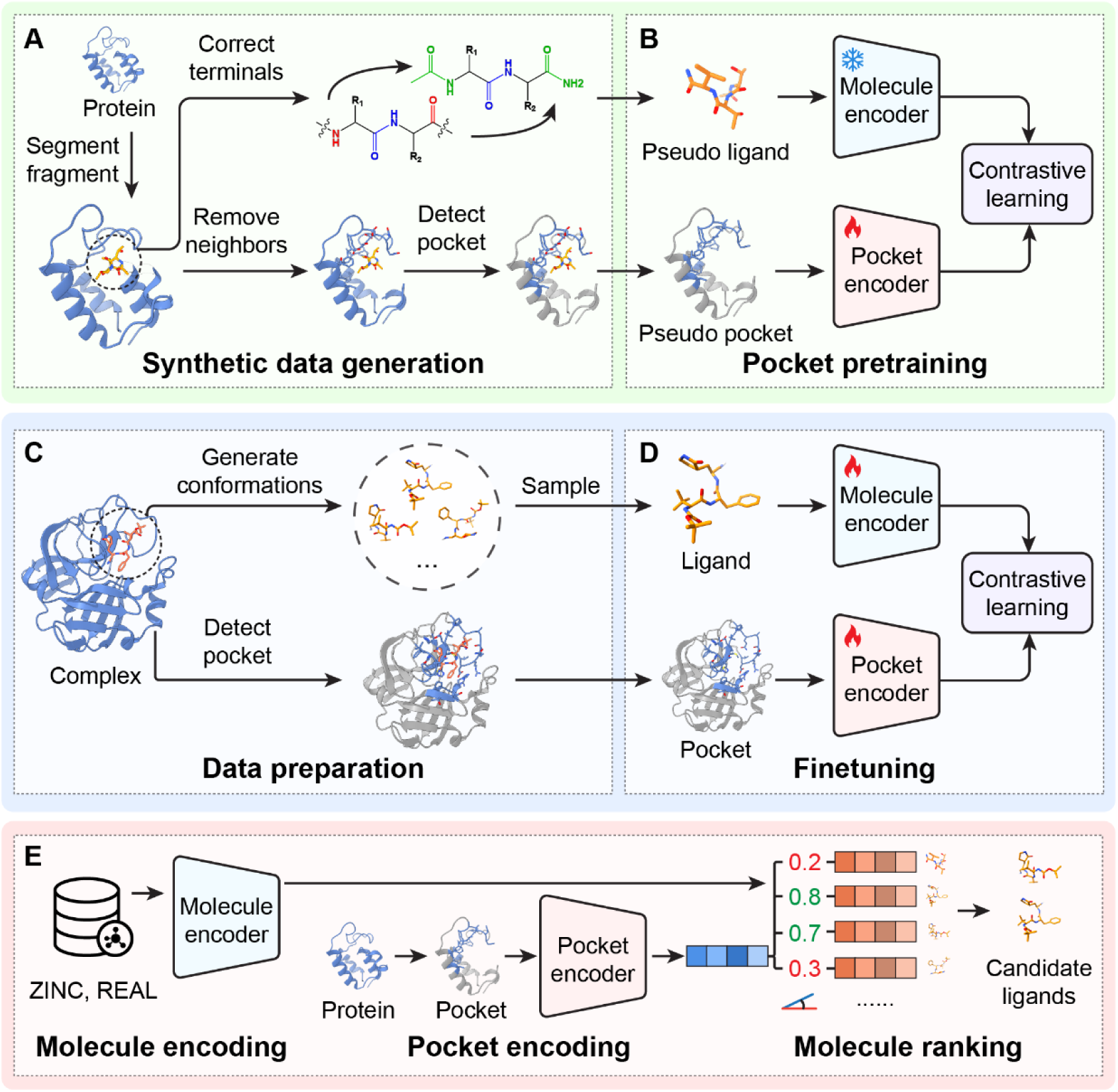
The framework of DrugCLIP. **(A)** In the pretraining stage, a large-scale synthetic dataset was created using the ProFSA strategy. Specifically, pseudo pocket-ligand pairs were constructed through a series of operations, including fragment segmentation, terminal correction, neighbor removal, and pocket detection, on protein data. **(B)** The pocket encoder is pretrained with pseudo pocket-ligand pairs in a contrastive distillation manner to transfer knowledge from a well-established molecular encoder to the pocket encoder. **(C)** During the fine-tuning process, experimentally determined protein-ligand pairs were used as training data, with multiple ligand conformations generated by RDKit. **(D)** In the fine-tuning stage, both the pocket and molecule encoders were updated using a contrastive loss, which maximizes the similarity between positive pairs and minimizes it between negative pairs. **(E)** The pipeline for virtual screening with DrugCLIP. The candidate molecules from the library were pre-encoded with the trained molecular encoder. For a given pocket, the trained pocket encoder converts it to a vector, and the cosine similarity is then utilized to select top ligands with the highest scores.

The training process of DrugCLIP includes two stages: pretraining and fine-tuning. The molecule and pocket encoders are pretrained with large-scale synthetic data and are further refined using experimentally determined protein-ligand complex structures during fine-tuning.

In the pretraining stage, the molecule encoder is initialized with Uni-Mol (*29*), a well-established molecule encoder. With the molecule encoder frozen, the pocket encoder is randomly initialized and trained to align with the molecule encoder using contrastive learning (Fig. 1B). We developed a Protein Fragment-Surrounding Alignment (ProFSA) framework (Fig. 1A) to generate large-scale synthetic data specifically tailored for contrastive pretraining. In this approach, short peptide fragments are extracted from protein-only structures to serve as pseudo-ligands, while their surrounding regions are designated as pseudo-pockets. Intra-protein interactions share many features with protein–ligand interactions, including hydrogen bonding, ionic attraction, π-π stacking, and other non-covalent interactions (Fig. S1). In previous research on ligand-binding protein design, intra-protein packing has also been exploited to determine statistically preferred orientations of chemical groups relative to the backbone of a contacting residue for protein-ligand interface modeling (*30*). This principle underlies the development of ProFSA. To further enhance model performance, we carefully calibrate the chemical property distributions of pseudo-ligands and binding pockets to closely match those observed in real complexes (Fig. S2 and S3), thereby minimizing the distribution gap between synthetic and real-world data. Technical details are provided in the “*The Pretraining of the Pocket Encoder*” section of the *Methods*.

Applying the ProFSA framework to PDB (*31*) data yielded 5.5 million pseudo-pocket and ligand pairs to facilitate the pretraining. The trained pocket encoder has been evaluated across various downstream tasks such as pocket property prediction (Table S1), pocket matching (Table S2), and protein-ligand affinity prediction (Table S3). Experimental results demonstrate that our pretrained pocket encoder exhibits strong performance, even in a zero-shot setting, outperforming many supervised learning-based models as well as physical and knowledge-based models. These results underscore the success of the pretraining stage in obtaining meaningful pocket representations.

After pretraining, the molecule and pocket encoders are further fine-tuned (Fig. 1D) using 40,000 experimentally determined protein-ligand complex structures collected by the BioLip2 database (*32*). Given that the binding conformations of molecules are unknown and only their topologies are provided in virtual screening, we implemented a random conformation sampling strategy for data augmentation by using RDKit (*33*) for conformation generation. This augmentation allows DrugCLIP to train on data that more accurately reflects the variability of real-world screenings, thereby enhancing the model’s performance and generalization ability.

In the screening process (Fig. 1E), we first use our trained encoders to represent molecules and pockets as vectors. Cosine similarities between the pocket and molecule embeddings are then computed, and candidate molecules are ranked according to these similarity scores. Since the molecule representations can be computed offline, DrugCLIP screening is highly efficient, requiring only the calculation of a simple cosine similarity and subsequent ranking. Therefore, with proper pre-encoding and parallelization, DrugCLIP can evaluate trillion-level target-molecule pairs with a single GPU accelerator, which is more than 10,000,000 times faster compared with traditional computational methods like molecular docking.

### Evaluating DrugCLIP performance with benchmarks and wet-lab experiments

We benchmarked DrugCLIP on two widely used virtual screening datasets, DUD-E (*20*) and LIT-PCBA (*21*). The DUD-E dataset contains 22,886 active compounds of 102 protein targets. For each active compound, 50 decoys with similar physical properties but different structures are generated. In contrast, LIT-PCBA comprises approximately 8,000 active and 2.64 million inactive compounds across 15 targets, derived from experimental results of the PubChem BioAssay database. DrugCLIP was compared with established physical-informed docking software, including Glide-SP (*5*), Autodock Vina (*6*), Surflex (*34*), and regression-oriented machine learning models, including NNscore (*13*), RFscore (*35*), Pafnucy (*36*), OnionNet (*12*), PLANET (*11*), Gnina (*37*), BigBind (*38*). In both sets of results (Fig. 2A and 2B, Table S4 and S5), DrugCLIP demonstrated a superior performance over all baseline methods in terms of EF1%, measuring the recall capacity of virtual screening models.

**Fig. 2.**
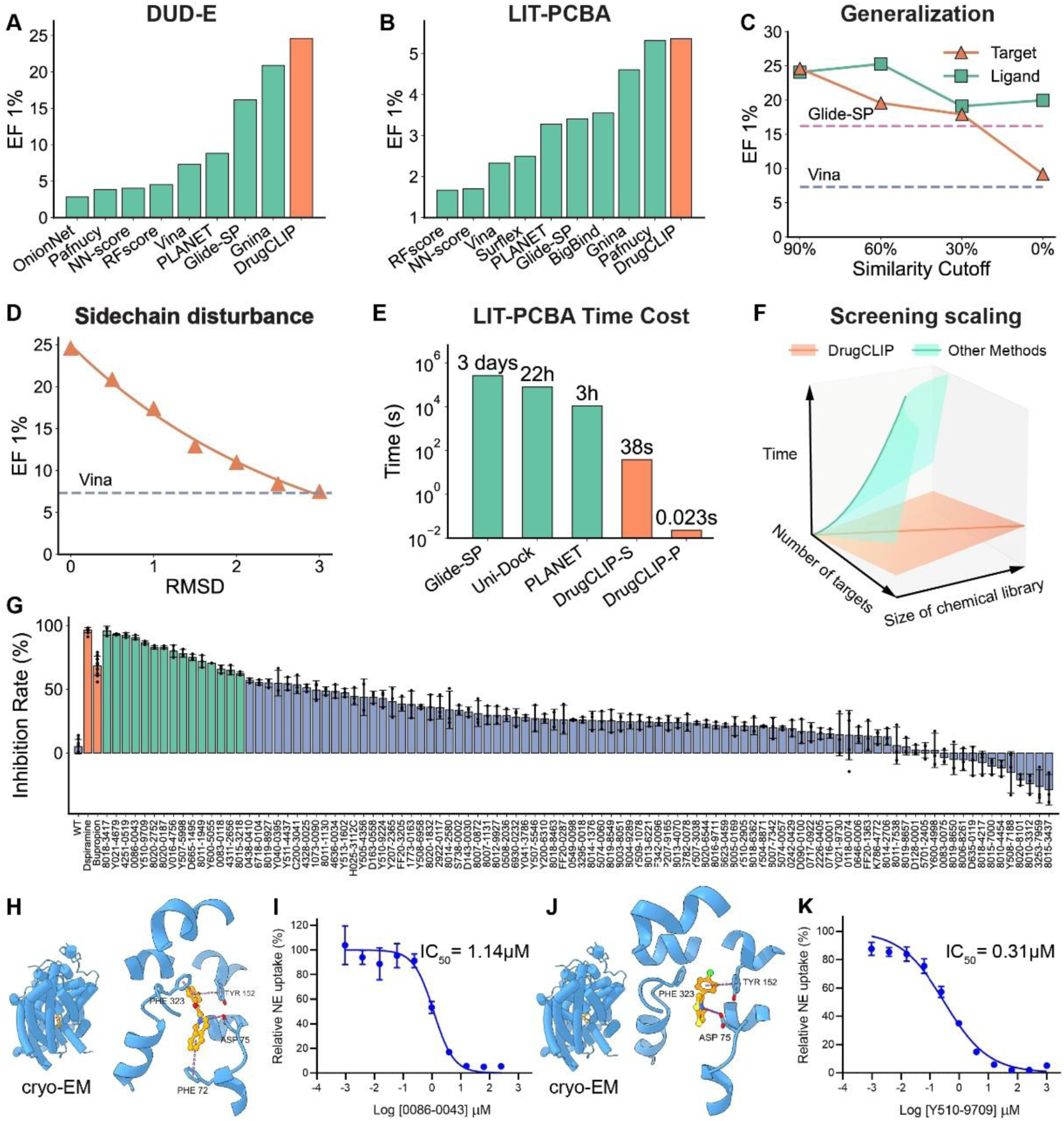
*In silico* benchmarking results of DrugCLIP and the wet-lab validation with NET. **(A)** The evaluation of DrugCLIP on the DUD-E dataset using the EF1% to assess model performance. The results of baseline models are taken from previous studies (*11, 48, 49*). **(B)** The evaluation of DrugCLIP on the LIT-PCBA dataset, also using the EF1% for performance measurement. The results of baseline models are taken from previous studies (*11, 21, 38, 49–51*). **(C)** The assessment of DrugCLIP’s generalization ability was conducted by varying the identity cutoffs between testing targets or molecules and training data in DUD-E, with Glide-SP and Vina represented as dashed lines. Protein similarities of 30%, 60%, and 90% are calculated by MMSeqs2 (*52*), and 0% indicates a protein family removal with HMMER (*53*) and PFAM (*54*). Molecular similarities of 30%, 60%, and 90% are calculated by Morgan2 (ECFP4) fingerprints (*55*), and 0% indicates a molecule series removal defined by generic Murcko scaffolds (*56*). **(D)** The evaluation of DrugCLIP’s robustness regarding errors in pocket side-chain conformations was conducted by using RMSD values ranging from 0 Å to 3 Å, with Vina shown as a dashed line for reference. **(E)** The screening speed on the LIT-PCBA dataset, compared with docking methods like Glide-SP and Uni-Dock, and the machine learning model PLANET. Speeds of baseline methods are taken from previous studies (*8, 11*). The time cost of Glide-SP is converted by using 128 CPU cores, as the setting of 16 CPU cores used in the original research is unfair to be compared with modern GPUs. For Uni-Dock, the time cost is estimated as 0.04s per ligand with 8 GPUs. As for DrugCLIP, sequential computing (DrugCLIP-S) of all LIT-PCBA targets on an A100 GPU will take 38 seconds, because the number of molecules and pockets in this dataset is too small to be properly parallelized on modern GPUs. Therefore, we also report a speed of parallel computing (DrugCLIP-P) by screening 10M molecules for 100k pockets, which will take around 25 minutes with an A100 GPU. Under this setting, it will only take 0.023 seconds for the same amount of computation as LIT-PCBA. **(F)** An illustration of time consumption as the screening scale increases, with the x-axis representing the size of the compounds library, the y-axis representing the number of targets, and the z-axis representing the time cost of virtual screenings. DrugCLIP (the orange line) has a computational complexity of *O*(M+N), where M is the number of targets and N is the number of compounds, whereas most existing methods (the green line) have a complexity of *O*(MN). **(G)** The evaluation of 100 DrugCLIP identified compounds with radio-ligand transportation assays for NET inhibitor at a concentration of 10 μM, and 15 compounds showed inhibition larger than 60%. **(H)** The complex structure of 0086-0043 and NET was determined with Cryo-EM. **(I)** The dose response curve of 0086-0043 in the radio-ligand transportation assay. **(J)** The complex structure of Y510-9709 and NET was determined with Cryo-EM. **(K)** The dose response curve of Y510-9709 in the radio-ligand transportation assay.

We also investigated the influence of molecule similarity, homology information, and protein structure accuracy on DrugCLIP’s performance. After removing training samples containing similar molecular substructures or scaffolds to the test set, the performance drop of DrugCLIP remains marginal. Notably, it consistently outperforms the widely used commercial virtual screening software Glide-SP (Fig. 2C, Table S6). The robustness of DrugCLIP is not only to unseen molecular structures, but also to new protein families. Remarkably, even when test protein families were entirely excluded from the training set, DrugCLIP still outperformed one of the most popular virtual screening methods AutoDock Vina (Fig. 2C, Table S7), highlighting its strong generalization capability to new targets. Moreover, DrugCLIP shows exceptional robustness by outperforming AutoDock Vina even with a 3 Å RMSD error in the side chain conformations of protein pockets (Fig. 2D), indicating its robustness to structural inaccuracies.

Furthermore, DrugCLIP is exceptionally efficient (Fig. 2E), making it highly suitable for large-scale screening tasks. For instance, DrugCLIP can complete the screening for LIT-PCBA in merely 38 seconds in the sequential computing mode, significantly faster than Glide docking (3 days), Uni-Dock (22 hours) (*8*), and another machine learning method PLANET (3 hours) (*11*). When a large number of molecules and pockets are evaluated, efficient parallel computing with GPUs can further reduce the time cost of the same amount of computation to 0.023 seconds. Moreover, the time consumption of DrugCLIP screening scales linearly with the simultaneous increase of target and molecule numbers (Fig. 2F), which can facilitate multi-target virtual screening.

These *in silico* results confirm that DrugCLIP possesses superior virtual screening capabilities, combining high performance, generalizability, robustness, and efficiency. In addition to *in silico* evaluation, we tested the DrugCLIP model on real-world targets using wet-lab experiments. We focused on two well-established targets for psychiatric diseases: the serotonin receptor 2A (5HT_2A_R) and the norepinephrine transporter (NET).

5HT_2A_R is an emerging target for antidepressant development. Its agonists have demonstrated strong and long-lasting antidepressant effects in both rodent models and humans (*39, 40*). Previous research suggests that the recruitment of β-arrestin2 following 5HT_2A_R activation is a key biochemical mechanism underlying these antidepressant effects (*41, 42*).

In a pilot virtual screening experiment, 78 top-ranked compounds were ordered from ChemDiv, Inc. (https://www.chemdiv.com/), which is also the supplier for the screening of another two targets in the following sections. Eight of the 78 compounds were identified as positive agonists in a calcium flux assay, exhibiting a minimal activity of 10% compared to serotonin (Fig. S4). The affinities of these compounds to 5HT_2A_R were further assessed using [³H]-labeled ketanserin competitive binding assays, with six showing a *K*_i_ of less than 10 μM (Table S8, Fig. S5 and S6). We then evaluated the cellular function of these hit compounds using NanoBit assays for β-arrestin2 recruitment, and all 6 compounds achieved an EC_50_ of less than 1 μM (Table S8, Fig. S5 and S6). The best compound achieves an affinity of 21.0 nM and exhibits an EC_50_ of 60.3 nM with an E_max_ of 35.8% in the NanoBit assay.

Following the success of 5HT_2A_R, we targeted a well-established drug target, the norepinephrine transporter (NET), for depression and attention deficit hyperactivity disorder (ADHD). Although there are multiple FDA-approved inhibitors (*43*), the structures of NET with or without its inhibitors in complexes were not solved until 2024 (*44–46*). The closest protein structure in our dataset is the dopamine transporter from *Drosophila* (*47*), which shares less than 60% similarity with NET. Therefore, screening against NET provides a more challenging test of our model’s ability to generalize to structurally new targets.

For this target, we ultimately selected 100 compounds considering chemical novelty and diversity. We tested their inhibition of NET protein by measuring the transport of [³H]-labeled norepinephrine in NET-containing liposomes. Among these compounds, 15% of them exhibited more than 60% inhibition of NET, with 12 compounds demonstrating greater potency than the widely used antidepressant bupropion.

Unlike previous NET inhibitors that typically feature aliphatic nitrogen atoms capable of forming a salt bridge interaction with ASP75 of NET (*44–46*), our screening identified several hits with positively charged aromatic nitrogen atoms. Notably, two of these compounds, 0086-0043 and Y510-9709, demonstrated better IC_50_ (with values of 1.14 μM and 0.31 μM, respectively) than bupropion (1.5 μM). Structural determination of the complexes between these compounds and the NET protein revealed that the aromatic rings indeed form more favorable interactions with NET: the isoquinoline ring of 0086-0043 engages in a T-shaped π-π interaction with PHE72, and the thiazole ring of Y510-9709 likely interacts with surrounding aromatic side chains like PHE323 and TYR152. These findings highlight the potential of the DrugCLIP model to provide new chemical insights for drug discovery.

### Applying DrugCLIP to AlphaFold-predicted structures

After validating the DrugCLIP model through both *in silico* and wet-lab experiments, we apply it to computationally predicted protein structures. Recent breakthroughs in protein structure prediction—most notably the near-complete coverage of the human proteome by AlphaFold2 (*22, 23*)—have provided structural insights into many important drug targets lacking experimental data. This opens new avenues for structure-based drug discovery beyond the limits of experimentally determined structures.

Virtual screening using AlphaFold-predicted structures remains a topic of debate. The primary concern is that these predicted structures may lack the accuracy needed to replicate experimental conformations and effectively filter out inactive molecules (*57, 58*). Despite this, some studies have shown that virtual screening with AlphaFold-predicted structures can still yield reasonable results for certain targets (*59, 60*). Given the robustness of DrugCLIP to sidechain inaccuracies (Fig. 2D), we further assess the influence of predicted structure using a specialized DUD-E subset for virtual screening of AlphaFold predictions and *apo* structures (*57*). First, we observed that DrugCLIP is robust to the conformational variability inherent in AlphaFold2-predicted or *apo* structures, as long as the binding pockets are accurately defined through structural alignment with *holo* references (as shown in Exp. Pocket in Fig. 3B). For protein targets without homology structures, software like Fpocket (*25*) is usually used to identify potential pockets. In our experiments, using Fpocket outcomes resulted in a significant performance drop for DrugCLIP, with the EF1% value decreasing from 29.3% to 19.0% (Fig. 3B, Table S10), reflecting similar challenges observed with docking methods in both virtual screening (*57*) and conformation prediction (*58*).

**Fig. 3.**
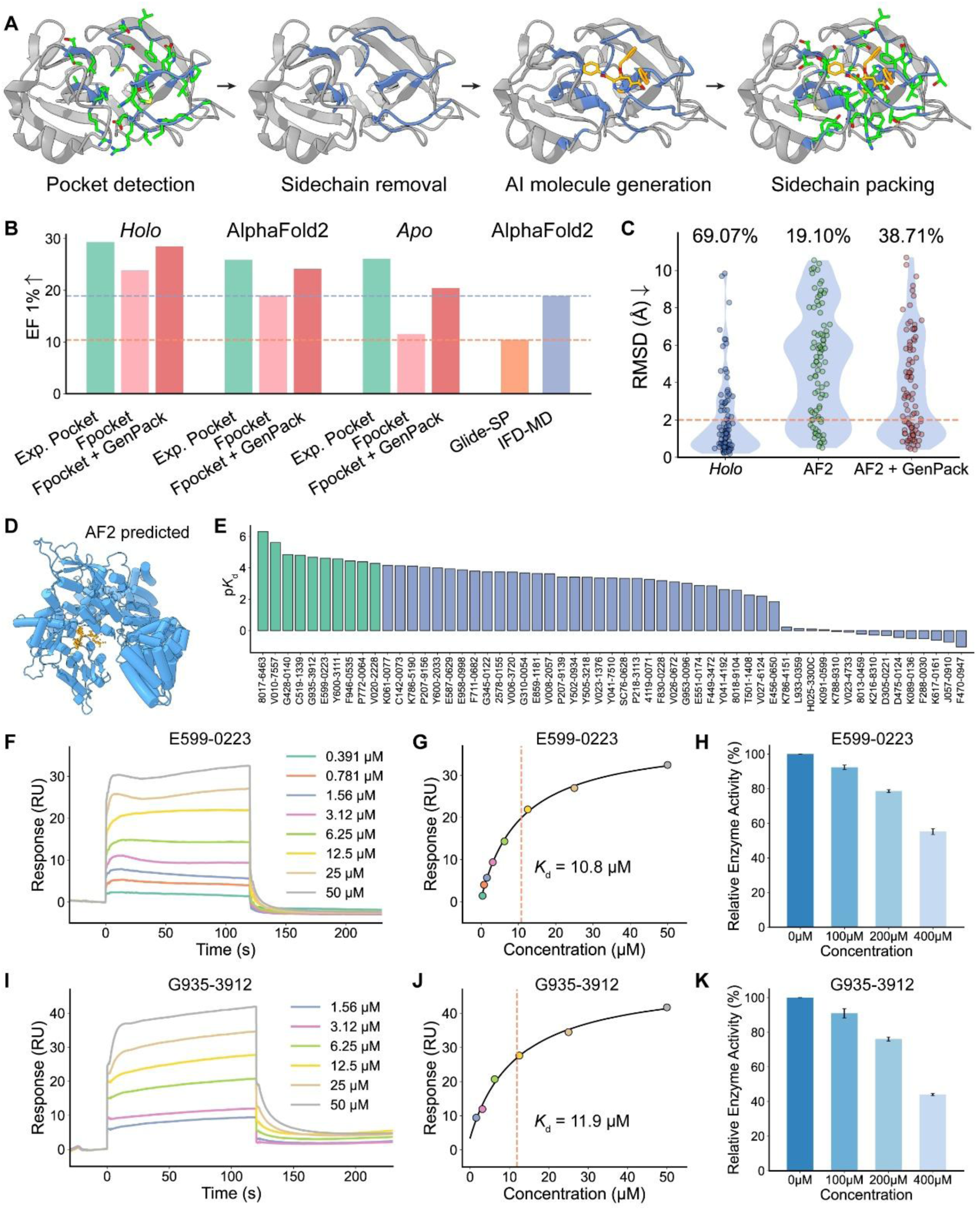
Applying DrugCLIP to AlphaFold-predicted structures with the aid of GenPack. (A) The GenPack (Generation-Packing) process for extracting pockets from AlphaFold2-predicted structures involves using Fpocket to detect initial pockets, removing sidechains, applying an AI-generative model to create molecules based on the backbone structure, and then performing sidechain packing with the generated molecules. (B) The EF1% comparisons for virtual screening on the DUD-E subset (*57*) of *holo*, AlphaFold2-predicted, and *apo* structures, using different pocket definitions: structural alignment to *holo* structures (Exp. Pocket), pockets detected by Fpocket (Fpocket), and pockets generated by GenPack (Fpocket + GenPack). The performances of Glide-SP and IFD-MD are given as references. (C) The redocking RMSD comparisons for different pocket definitions: *holo*-pocket, pockets on AlphaFold2-predicted structures, and pockets on AlphaFold2-predicted structures refined by GenPack. The orange dashed line indicates the RMSD threshold of 2 Å, and the corresponding docking success rates are labeled above each column. (D) AlphaFold2-predicted structure of TRIP12, and the pocket used for virtual screening with DrugCLIP (orange dots). (E) p*K*d values of 57 selected compounds measured by single-cycle SPR in initial screening; green color indicates hit compounds with their *K*d value lower than 50 μM, validated by following multi-cycle SPR assays. (F) Sensorgram of the multi-cycle SPR assay for E599-0223. (G) Steady-state binding curve of the multi-cycle SPR assay for E599-0223. (H) Enzyme activities of TRIP12 under different concentrations of E599-0223. (I) Sensorgram of the multi-cycle SPR assay for G935-3912. (J) Steady-state binding curve of the multi-cycle SPR assay for G935-3912. (K) Enzyme activities of TRIP12 under different concentrations of G935-3912.

To improve the utility of AlphaFold-predicted structures, we developed a strategy called GenPack (Generation-Packing, Fig. 3A). This strategy involves training molecular generative models conditioned on the backbone structures of protein pockets. While the generated molecules may not always be synthesizable, they help to localize pockets more precisely and induce the pocket conformation into a more suitable state. After this generation step, side chains are reintroduced, and the overall conformation is refined using physical force fields. With the GenPack strategy, we significantly enhanced the screening power of AlphaFold-predicted structures, increasing EF1% value on the DUD-E subset from 19.0% to 24.1% (Fig. 3B, Table S10). As for *apo* structures, the performance boost from GenPack is more significant, where EF1% was improved from 11.5% to 20.4% (Fig. 3B, Table S10). Compared to the previous state-of-the-art virtual screening method for *apo* or AlphaFold-predicted structures, IFD-MD (*57, 61*), our approach achieves superior performance in terms of active molecule enrichment. Additionally, GenPack improves the docking success rate when using AlphaFold2-predicted receptors, increasing it from 19.1% to 38.7% across all DUD-E targets with available AlphaFold2 structures (Fig. 3C, Table S12).

To further understand the mechanism of GenPack’s performance boost to DrugCLIP and molecular docking, we conducted additional experiments to evaluate the pocket refinement by GenPack.

We first investigated whether this process could refine pocket conformations to better resemble *holo* structures. Surprisingly, GenPack refinement did not improve the overall side-chain RMSD relative to *holo* structures. Furthermore, for AlphaFold2-predicted structures—regardless of whether GenPack refinement was applied—we observed no correlation between side-chain RMSD and either docking performance (measured by ligand docking pose RMSD, Fig. S10D) or screening performance (measured by ΔEF1%, Fig. S10B). Based on these findings, we conclude that GenPack does not improve the pocket conformation of AlphaFold2 structures, and pocket side-chain accuracy appears to have limited influence on virtual screening or docking performance in our setting. Similar results were also observed in the previous research of molecular docking with AlphFold2 predictions (*58*).

Since automated tools like Fpocket were less precise in detecting ligand-binding pockets compared to structural alignment approaches, we then conducted additional experiments to further investigate whether GenPack improves the pocket detection and localization for AlphaFold2 predictions. We found that the decrease in virtual screening performance, measured by ΔEF1%, is correlated with the precision of pocket detection, quantified by the intersection-over-union (IoU) between predicted and *holo* pockets (p < 0.005, Fig. S10A). Importantly, GenPack refinement improved the pocket IoU scores (the distribution curves on top of Fig. S10A), suggesting that it enhances pocket definition and, as a result, contributes to improved virtual screening outcomes. Nevertheless, the localization refinement is not correlated to the docking performance (Fig. S10C).

Taken together, these results demonstrate that DrugCLIP, with the aid of GenPack, achieves superior virtual screening performance on *apo* or AlphaFold2-predicted structures compared with physically informed methods like IFD-MD.

Beyond *in silico* evaluations, we further demonstrate the capabilities of GenPack and DrugCLIP using a novel and promising biological target, thyroid hormone receptor interactor 12 (TRIP12). TRIP12 is an E3 ubiquitin ligase (*62*) that represents a potential drug target implicated in cancers and neurodegenerative diseases. TRIP12 mediates the ubiquitination of p14ARF, leading to its degradation and consequently suppressing p53 activity in cancer cells (*63*). In the nervous system, TRIP12 functions as a key regulator of GCase (glucocerebrosidase), targeting it for ubiquitin-mediated degradation, which leads to α-synuclein accumulation and aggregation, a pathological hallmark of Parkinson’s disease (*64*). Despite its biological significance, TRIP12 remains challenging for drug discovery. Structures containing the catalytic HECT domain and small-molecule inhibitors for this target have not been released to date. This absence of structural data and chemical starting points positions TRIP12 as a particularly challenging yet scientifically valuable target for validating the generalization capabilities of DrugCLIP and GenPack.

We applied DrugCLIP to the predicted binding pocket near the catalytic site of TRIP12 (Fig. 3D), as identified from the AlphaFold-predicted structure. The top 1% of ranked compounds were finalized to a selection of 57 candidate compounds for experimental validation. Among these, 10 compounds demonstrated *K*_d_ values lower than 50 μM, as determined by surface plasmon resonance (SPR) assays, yielding a hit rate of 17.5% (Fig. 3E, Fig. S11, Table S14). The two best compounds, E599-0223 and G935-3912, showed affinities to TRIP12 of 10.8 μM and 11.9 μM, respectively (Fig. 3F, G, I, J). Additionally, their dose-dependent inhibition of TRIP12’s ubiquitination activity was confirmed using fluorescent ubiquitination assays (Fig. 3H and K, Fig. S12), and they showed no off-target inhibition to E1 ubiquitin-activating enzyme and E2 ubiquitin-conjugating enzyme at the highest concentration (Fig. S13). To the best of our knowledge, these compounds represent the first publicly reported inhibitors of the ubiquitination function of TRIP12.

Together, *in silico* and experimental results demonstrate that DrugCLIP is an effective virtual screening tool for AlphaFold-predicted protein structures. These findings highlight a promising path forward for structure-based drug discovery targeting proteins lacking experimentally determined structures.

### Genome-wide virtual screening with DrugCLIP

Finally, we introduced a genome-wide virtual screening pipeline to facilitate future drug discovery. We began with splitting all AlphaFold predictions of human proteins into high-confidence regions based on plDDT and PAE scores. For each region, we used homology alignment and Fpocket (*25*) along with GenPack to detect potential pockets. The DrugCLIP model was then employed to screen over 500 million drug-like molecules from the ZINC (*26, 27*) and Enamine REAL (*28*) databases. The screening process, which involved more than 10 trillion scoring operations on protein-ligand pairs, was completed in about 24 hours on a single computing node equipped with 8 A100 GPUs. The top-ranked molecules were then clustered and further evaluated using molecular docking, filtering out poor poses with Glide score > -6 kcal/mol. The final database contains over 2 million potential hit molecules for more than 20,000 pockets from 10,000 human targets. All molecules, docking scores, and poses have been made freely accessible at https://drug-the-whole-genome.yanyanlan.com (Fig. 4A), facilitating further research and drug discovery processes.

**Fig. 4.**
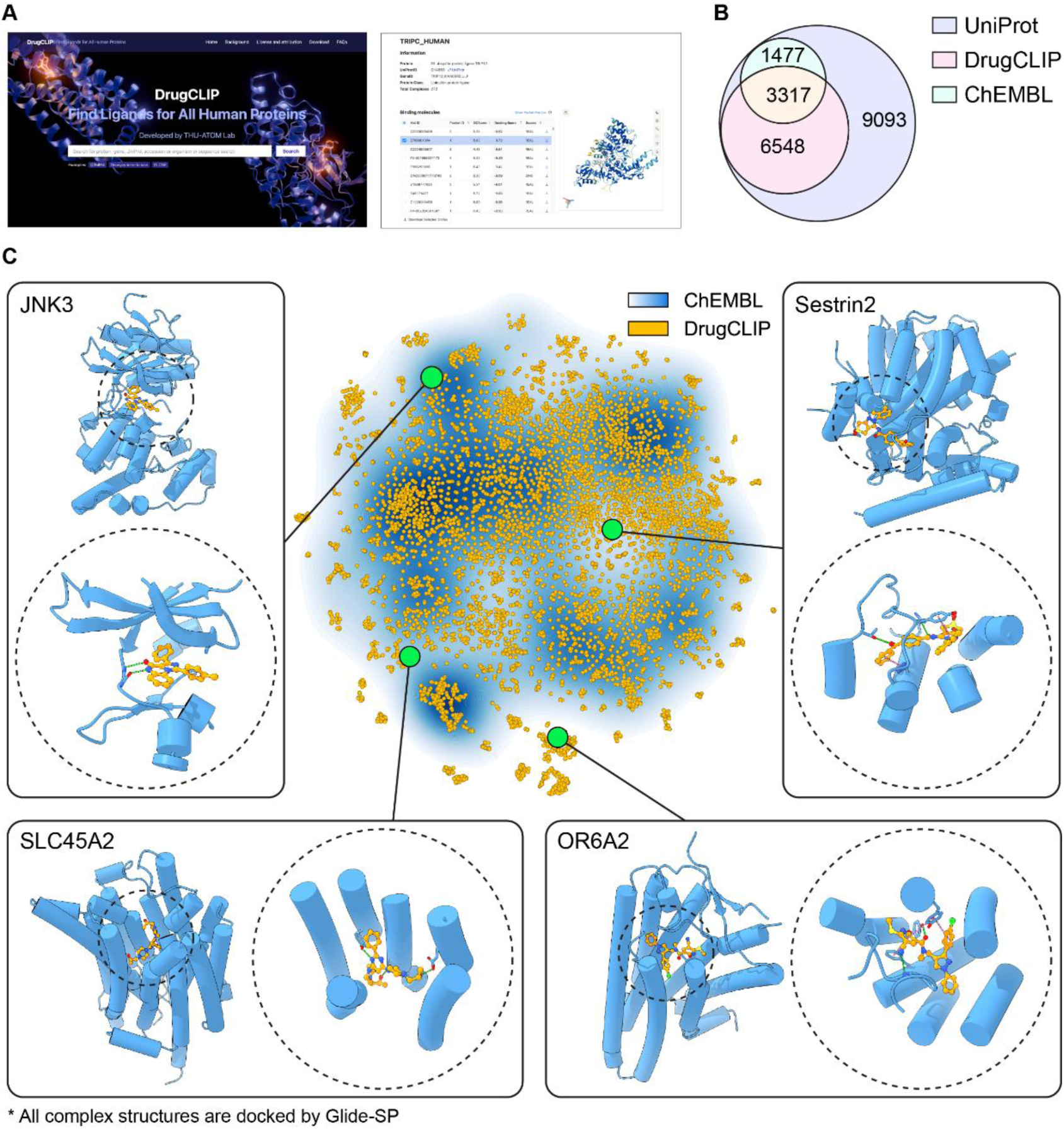
DrugCLIP enables genome-wide virtual screening. **(A)** The webpage for accessing our genome-wide virtual screening results at https://drug-the-whole-genome.yanyanlan.com **(B)** The Venn diagram of target numbers in different databases, with UniProt, DrugCLIP, and ChEMBL shown as different circles. **(C)** The t-SNE visualization and examples for the genome-wide virtual screening results. Yellow dots indicate targets in our database, while the blue-white gradient represents targets in the ChEMBL database, with density ranging from high (blue) to low (white).

Our genome-wide screening results cover a more extensive range of targets than ChEMBL (*65*), one of the most comprehensive databases for bioactive molecules. While UniProt (*1*) contains 20,436 reviewed human proteins, the latest ChEMBL release (ChEMBL 34) covers 4,810 of them. Moreover, not all targets in the ChEMBL database have high-affinity small-molecule binders; some targets only have peptide or antibody binders, or merely vague results from low-quality assays. In contrast, our database spans 9,908 targets, more than twice the number in ChEMBL and covers nearly half of the human genome (Fig. 4B). To visualize the difference between the two protein spaces, we encoded all protein sequences using the ESM1b model (*66*). The t-SNE plot shows that our space encompasses a broader range of proteins, including many that are not closely related to those in ChEMBL (Fig. 4C).

Our database includes a diverse range of targets, from well-studied proteins to less-explored members of well-known families, as well as proteins with limited pharmacological understanding (Fig. 4C). For example, the c-Jun N-terminal kinase 3 (JNK3) is a classical kinase target with many ligand-bound crystal structures (*67, 68*). DrugCLIP identified molecules that bind to the ATP-binding pockets, forming H-bonds with backbone atoms of MET149 in the hinge region. SLC45A2 belongs to the solute carrier (SLC) superfamily, many of which are important drug targets. Nevertheless, SLC45A2 has limited pharmacological studies. This gene plays a crucial role in pigmentation (*69*) and is widely expressed in cutaneous melanomas (*70*), with evidence suggesting its oncogenic potential (*71*). All molecules in the database could bind near L374, which is an important site for protein stability (*69*), thus having potential modulatory effects. Another interesting example OR6A2 belongs to the olfactory receptor family, whose members are mainly found to be expressed in olfactory receptor neurons, yet many of them are expressed in various other tissues with unexplored pharmaceutical potentials (*72*). OR6A2 is expressed in macrophages, sensing blood octanal and promoting the formation of atherosclerotic plaques (*73*). Our predicted molecules fit the orthosteric pocket of OR6A2 and can serve as potential inhibitors for treating atherosclerosis. The final example Sestrin-2 can sense leucine (*74*) and promote drug resistance of cancer cells (*75*), which belongs to a unique highly-conserved stress-inducible protein family (PF04636 or IPR006730) with only three members in the human genome. Our database contains predicted molecules that bind to the same pocket of leucine (*76*) that may serve as good starting points for anti-cancer therapies. These examples highlight the potential of our database as a valuable resource for exploring the undrugged genome and facilitate future drug discovery.

## Conclusions and Discussions

With the rapid advancement of protein structure prediction methods and the availability of a comprehensive atlas of predicted protein structures for human and disease-related species (*23, 77*), we have entered a new era where effective drug discovery for all disease-related targets is within reach. In this paper, we introduce DrugCLIP, a groundbreaking contrastive learning based virtual screening approach that aims to achieve genome-wide drug discovery. The efficacy of DrugCLIP has been rigorously validated through both *in silico* benchmarks and wet-lab experiments. In well-established benchmarks, DrugCLIP consistently outperformed traditional docking software and contemporary machine learning models. Notably, for the 5HT_2A_R and NET targets, DrugCLIP identified diverse high-affinity binders and novel chemical entities. We further validated the capability of DrugCLIP on TRIP12, a particularly challenging target with no available structural and chemical information. DrugCLIP has identified the first reported small-molecule inhibitors of TRIP12, providing valuable starting points for this promising therapeutic target. These findings underscore the potential of DrugCLIP model as a reliable tool for virtual screening in real-world drug development. We demonstrate its application through a genome-wide virtual screening campaign, encompassing more than 20,000 pockets across approximately 10,000 human proteins, using a chemical library of 500 million molecules from ZINC and Enamine REAL. Remarkably, DrugCLIP completes this trillion-level virtual screening campaign in just 24 hours using just a single computational node with 8 GPU accelerators. Beyond the screening results, we have generated over 2 million high-confidence protein-ligand complex structures accompanied with their docking score. By making this extensive database freely accessible, we aim to make a substantial contribution to the research community, accelerating drug discovery and fostering innovation in therapeutic development.

DrugCLIP is more than just a new tool. It represents a transformative shift in the development of new therapeutics, heralding a new paradigm in drug discovery. Its genome-wide virtual screening capability opens the door to truly end-to-end drug discovery on a genomic scale, allowing researchers to screen all relevant targets simultaneously, rather than focusing on a few promising targets. This expansive approach facilitates the creation of customized chemical libraries for advanced phenotypic screening with high-fidelity models such as organoids (*78–80*) or humanized mice (*81–83*), potentially reducing failure rates in drug development.

DrugCLIP paves the way for new advancements in AI-driven drug discovery. Its outstanding efficiency allows the screening scale to the largest ultra-large chemical library available today, e.g., 48 billion-compound Enamine REAL Space library. This effort pushes the boundaries of what virtual screening can achieve in drug discovery. Moreover, the release of these genome-wide virtual screening results could serve as a valuable resource for molecular generation, particularly through a retrieval-augmented generation approach (*84, 85*), enhancing our capacity for drug discovery and design.

## Acknowledgement

We would like to take this opportunity to thank Dr. Xiaobing Deng, Prof. Zhenming Liu, Dr. Qiwei Ye and Bo Qiang for their valuable discussions and insightful comments. The manuscript is polished and revised with ChatGPT-3.5 and ChatGPT-4. IHCH-7079 is a gift from Prof. Sheng Wang and Prof. Jianjun Cheng, and we sincerely thank their generous help. Some wet-lab experiments are conducted with the aid of the Center of Pharmaceutical Technology at Tsinghua University, and we sincerely appreciate their help. This work is supported by the National Key R&D Program of China (No.2021YFF1201600 to Y.L. and No.2020YFA0509301 to C.Y.), the National Natural Science Foundation of China (32341016 and 32171204 to C.Y.; 22137005 and T2488301 to L.L.), Beijing Academy of Artificial Intelligence, and Beijing Frontier Research Center for Biological Structure Fundings.

## Supplementary Results

### Benchmarking the performance of pocket pretraining with ProFSA

To test the performance of the pretrained pocket encoder, we benchmark the encoder on three major benchmarks. The first task is about the pocket druggability prediction. We assess the effectiveness of ProFSA in predicting various physical and pharmaceutical properties of protein pockets, utilizing the druggability prediction dataset created by Uni-Mol [1]. This dataset comprises four separate regression tasks: Fpocket score, Druggability score, Total Solvent Accessible Surface Area (SASA), and Hydrophobicity score. The evaluation metric employed for these tasks is the Root Mean Square Error (RMSE), which measures the accuracy of the predictions. The baseline model we compared is the pocket encoder from the Uni-Mol [1]. The result is shown in **Table S1**.

The second task is the zero-shot pocket matching, for which we use two datasets: the Kahraman dataset [2] and the TOUGH-M1 dataset [3]. The Kahraman dataset contains matched pockets from two non-homologous proteins that bind to the same ligand. It consists of 100 proteins binding to 9 different ligands. We use a reduced version of this dataset, excluding 20 PO_4_ binding pockets due to their low number of interactions. The TOUGH-M1 dataset, on the other hand, involves relaxing identical ligands to identify similar pockets and comprises 505,116 positive and 556,810 negative protein pocket pairs derived from 7,524 protein structures. The baseline models we employed encompass various approaches, including PocketMatch [4], DeeplyTough [5] and IsoMIF [6]. Additionally, we consider established software tools like SiteEngine [7] and TM-align [8]. We also incorporate pretraining strategies, such as Uni-Mol [1] and CoSP [9]. The result is shown in **Table S2**.

The third task is binding affinity prediction. We use the widely recognized PDBBind dataset (v2019) for predicting ligand binding affinity (LBA), following the strict 30% or 60% protein sequence identity splits and preprocessing protocols specified by Atom3D. These strict data splits are crucial for providing reliable and meaningful comparisons, especially in evaluating the robustness and generalization capabilities of the models. For each protein-ligand pair, we concatenate the protein embedding from our pretrained pocket encoder with the molecular embedding from the Uni-Mol molecular encoder and pass this combined representation through a multilayer perceptron (MLP) to generate the final binding affinity prediction. For our baseline models, we utilize a diverse range of methods including DeepDTA [10], B&B [11], TAPE [12], ProtTrans [13], HoloProt [14], IEConv [15], MaSIF [16], and several ATOM3D variants—3DCNN, ENN, and GNN [17]. Additionally, we incorporate ProNet [18] and pretraining approaches such as GeoSSL [19], EGNN-PLM [20], DeepAffinity [21], and Uni-Mol [1]. The result is shown in **Table S3**.

## Data and Code availability

All input data are freely available from public sources.

For ProFSA pretraining, the PDB database can be acquired from https://www.wwpdb.org/ftp/pdb-ftp-sites. The processed dataset is available at HuggingFace: https://huggingface.co/datasets/THU-ATOM/ProFSADB. Related code and model weights are available at: https://github.com/THU-ATOM/ProFSA.

DrugCLIP is fine-tuned using the BioLip2 dataset, available on: https://zhanggroup.org/BioLiP/index.cgi. For the 6-fold version, please refer to **Supplementary Materials 1**. For all similarity-based splits, refer to **Supplementary Materials 2** for the list of pre-filtered PDB IDs. Related code and model weights are available at: https://github.com/bowen-gao/DrugCLIP.

GenPack is trained using the PDBBind2020 dataset, available at: https://www.pdbbind-plus.org.cn/download. For the list of pre-filtered PDB IDs based on pocket similarity to DUD-E, please refer to **Supplementary Materials 3**. Related code and model weights are available at: https://github.com/THU-ATOM/Pocket-Detection-of-DTWG. Datasets for benchmarking are downloaded from their official websites, including DUD-E (https://dude.docking.org/), LIT-PCBA (https://drugdesign.unistra.fr/LIT-PCBA/), and ATOM3D (https://www.atom3d.ai/). For the subset of 27 DUD-E targets for *apo* and AlphaFold predictions, please refer to its original publication [22]. For all 96 DUD-E targets with available AlphaFold2 predictions, please see **Supplementary Materials 4** for their gene names. The pocket matching and pocket property prediction benchmarks are acquired from their original publications [1, 2, 3]. For wet-lab validation, we provide a reference pipeline using DrugCLIP and molecular docking. Note that human evaluation of candidate molecules can influence virtual screening outcomes. The reference pipeline is available at: https://github.com/THU-ATOM/DrugCLIP_screen_pipeline.

All docking poses from the genome-wide screening are available at: https://drug-the-whole-genome.yanyanlan.com/. The unfiltered data can be accessed at: https://huggingface.co/datasets/THU-ATOM/GenomeScreen.

## Materials and Methods

### The design of DrugCLIP

The DrugCLIP model has a molecule encoder and a pocket encoder. These two encoders are aligned by contrastive learning.

Both encoders are based on the Uni-Mol architecture [1], a transformer architecture that takes 3D atomic features as input. For the molecule encoder, we directly utilize the pretrained weights from Uni-Mol for initialization, leveraging its learned representations for small molecules. The pocket encoder is pretrained to be aligned with the molecule encoder in a contrastive distillation manner [23] with the ProFSA dataset.

The training of the DrugCLIP model is under a contrastive learning framework. Given a batch of encoded protein-ligand pairs {(*p*_1_, *m*_1_), (*p*_2_, *m*_2_), . . ., (*p*_*n*_, *m*_*n*_)}, where *p*_*i*_ is the embedding of the protein pocket *i* obtained from the pocket encoder. *m*_*i*_ is the embedding of the corresponding ligand *i* encoded by the molecular encoder.

The objective is to learn embeddings such that the representations of true (positive) protein-ligand pairs are closer together in the embedding space, while the representations of incorrect (negative) pairs are further apart.

To accomplish this, we use a contrastive learning framework with a batch softmax approach, which involves two main loss functions.

The first loss is designed to find the correct ligand *m*_*i*_ for a given protein pocket *p*_*i*_. The loss function for this objective can be written as:

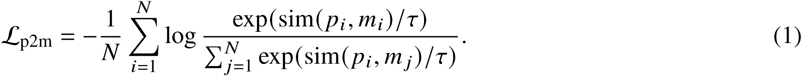

sim(*p*_*i*_, *m* _*j*_) represents a similarity measure between the protein pocket embedding *p*_*i*_ and ligand embedding *m* _*j*_. Here we use the cosine similarity. τ is the temperature parameter controlling the sharpness of the softmax distribution.

The second loss aims to find the correct protein pocket 73 *p_i_* from a batch of pocket candidates given a ligand *m_i_*:

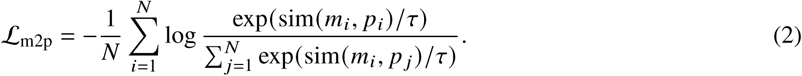

The final contrastive loss for training the model is the sum of the two losses:

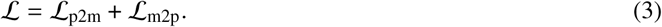

### The pretraining of the pocket encoder

The pocket pretraining uses the protein fragment-surroundings alignment (ProFSA) framework. The Protein Data Bank (PDB) [24] contains a vast amount of protein-only data. Interestingly, small molecule-protein interactions often mirror the non-covalent interactions found within proteins themselves [25]. Such similarity is shown in **Fig. S1**. Leveraging this similarity, we first extract fragments from protein structures that closely resemble known ligands and define the surrounding regions as the associated pockets of these pseudo-ligands.

In the initial phase, we iteratively isolate protein fragments ranging from 1 to 8 residues, ensuring these segments are continuous from the N-terminal to the C-terminal while excluding any discontinuous sites or non-standard amino acids. To minimize artifacts introduced by the cleavage of peptide bonds during fragment segmentation, we apply terminal modifications: acetylation at the N-terminus and amidation at the C-terminus. For the N-terminus, we cap with an acetyl group constructed from the actual C, CA, and O atoms of the previous residue in the protein structure. For the C-terminus, we apply amidation using the N atom from the following residue. All capping atoms are extracted directly from neighboring residues within the same experimentally resolved structure, ensuring physical plausibility and avoiding steric clashes. These modifications result in the formation of pseudo-ligands.

In the subsequent phase, to focus on long-range interactions, we exclude the five nearest residues on each side of the fragment. We then designate the pocket as the surrounding residues that have at least one heavy atom within a 6 Å distance from the fragment.

The derived pseudo-complexes undergo stratified sampling based on the distribution observed in the PDBbind2020 dataset [26, 27], considering critical parameters such as pocket sizes (measured by the number of residues) and ligand sizes (expressed as effective residue numbers, calculated by dividing the molecular weight by 110 Da). Another key metric is the relative solvent-accessible surface area (rBSA), which we calculate using the FreeSASA package [28]. The pseudo-complexes are sampled to approximate the distributions seen in the PDBbind dataset [26, 27], particularly in terms of rBSA and the joint distribution of pocket-ligand size. This ensures the dataset’s representativeness and its suitability for training ligand-oriented contrastive learning models, as shown in **Fig. S2** and **Fig. S3**.

The final dataset comprises 5.5 million ligand-protein pairs, significantly larger than any existing protein-ligand complex structure dataset.

The ProFSA pretraining objective is also a batch softmax loss, where the Uni-Mol molecular encoder is used for the pseudo-ligands. During the training, the weights of the molecular encoder are frozen. This setup allows us to distill knowledge from the pretrained molecular encoder into the pocket encoder, enhancing its ability to learn interaction-aware representations of protein pockets. During the pretraining phase, the batch size is 4 × 48 on 4 NVIDIA A100 GPUs. We use the Adam optimizer with a learning rate of 0.0001. The max training epochs is 100. We use polynomial decay for the learning rate with a warmup ratio of 0.06.

### The fine-tuning process of DrugCLIP

We use ligand-receptor complex data from the BioLip2 [29] database, removing redundant entries (proteins with a sequence identity > 90% and binding to the same ligand) and cleaning the dataset to obtain around 43,980 high-quality protein-ligand complexes (a list of all PDB IDs in the training set is included the **Supplementary Materials 1**). The binding pocket for each protein is defined as the set of residues with at least one atom within 6 Å of any ligand atom. During training, we use ligand conformations sampled by RDKit rather than their co-crystal conformations to minimize the discrepancy between training and actual virtual screening conditions, as the true conformations of candidate molecules are unknown during screening. This approach reflects the practical scenario of virtual screening, where true crystal conformations are typically unavailable for large compound libraries. To enhance model robustness, we apply a data augmentation strategy by generating up to 10 conformations per molecule. In each training epoch, one conformation is randomly selected, allowing the model to learn from structural variability and generalize better across different conformations.

We use an ensemble model for most applications unless stated otherwise, including wet-lab validations with the NET and TRIP12 target and the final genome-wide virtual screening. These applications follow a 6-fold cross-validation strategy: the dataset is split into six folds, and the model is trained on five while validated on the remaining fold in each iteration.

For the 5HT_2A_R target, we adopt an 8-fold cross-validation strategy and apply data augmentation techniques, including HomoAug and ligand augmentation using the ChEMBL dataset [30], following the DrugCLIP method [31].

We train the model with a batch size of 48 on 4 NVIDIA A100 GPUs. The optimizer is Adam with a learning rate of 1e-3. adam betas are 0.9 and 0.999, adam eps is 1e-8. The max epochs is set to be 200. We use polynomial decay for the learning rate and the warm-up ratio is 0.06.

### Ensembling multiple pockets and models during screening

As described above, we obtain six model weights through 6-fold cross-validation. During virtual screening, these six model weights are used to generate six different predictions, which are then combined using mean pooling to achieve a robust virtual screening result.

During virtual screening, a target of interest may have multiple pocket conformations. For any candidate molecule, we use a max pooling approach to determine the maximum score between the molecule and the different pockets. However, because different pockets may have varying score ranges, this can introduce bias when applying max pooling. To address this, we normalize the scores using an adjusted robust z-score before performing the max pooling. Specifically, for a list of scores *X*:

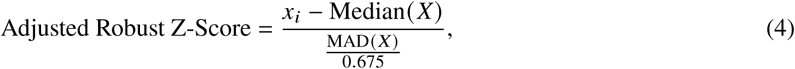

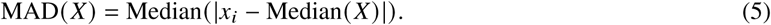

### *In silico* validation with DUD-E and LIT-PCBA dataset

The DUD-E (Directory of Useful Decoys: Enhanced) dataset [32] is a widely used resource in drug discovery research, particularly for evaluating the performance of virtual screening methods. It includes data on 102 protein targets with 22,886 active compounds known to bind to these proteins, along with a set of decoy molecules that are similar in physical properties but different in structure from the active compounds.

LIT-PCBA [33] is a benchmark dataset derived from the PubChem BioAssay database, designed for evaluating machine learning models in virtual screening and drug discovery. In the LIT-PCBA dataset, actives and decoys are defined based on experimental results from the PubChem BioAssay database. The dataset contains approximately 1.5 million compounds across 15 targets.

For the DUD-E and LIT-PCBA benchmarks, we use a single (non-ensemble) model trained on datasets filtered at 90% sequence identity using MMseqs2 [34]. In the homology removal test on the DUD-E benchmark, a single model is trained and evaluated on datasets filtered at 30%, 60%, and 90% identity via MMseqs2. The most stringent homology removal is performed using HMMER [35, 36] and the Pfam database [37]. As for ligand novelty analysis, we excluded training samples that their molecules are similar to any active molecules in the DUD-E test set by ECFP4 (Morgan2 by RDKit) similarity at cut-offs of 30%, 60% and 90%. For the strictest test, we remove all training samples that share the same generic Murcko scaffold as active molecules in DUD-E (indicated by 0% similarity in **Fig. 2C**.

For each target in the DUD-E or LIT-PCBA dataset, we rank candidate molecules (including both actives and decoys) based on their cosine similarity score. This score is calculated between the encoded embeddings of the pocket and molecule using the DrugCLIP model. The Enrichment Factor (EF) is then calculated to evaluate the ability of the model to prioritize active compounds over decoys. EF quantifies how many more actives are retrieved within the top-ranked subset than would be expected by random chance. It is typically defined as:

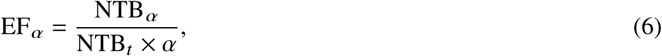

where NTB_α_ is the number of true active compounds (True Binders) identified within the top α fraction of the screened list. NTB_*t*_ is the total number of true active compounds in the entire dataset. α is the fraction of the dataset considered. In this manuscript, we use α = 1%, denoted as EF1%.

EF is closely related to the concept of recall capacity in the early retrieval stage. Specifically, recall at the top α fraction is defined as Recall_α_ = <u>^NTBα^</u>. Substituting this into the EF formula yields:

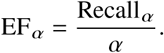

This shows that EF_α_ is essentially a normalized form of early recall, indicating how much better the model performs compared to random selection. A higher EF implies a stronger early recall capacity — the ability to identify true actives within the top-ranked results when only a small portion of the dataset is considered.

### Molecule selection for wet-lab experiments of 5HT_2A_R, NET and TRIP12

In general, for each target, DrugCLIP automatically enriches 1% to 2% molecules of the given chemical library. Around 200 chemically diversified molecules were picked from the top-ranked molecules by human experts, with the aid of clustering software and fingerprints like MACCS or ECFP. Glide docking will be performed on at most these picked diversity sets, and all molecules with docking scores lower than -6 will be manually examined. Based on the chemical structures, docking poses, and docking scores, around 100 molecules will be ordered from the chemical supplier. Additional physical property filters and novelty filters will be applied if necessary.

The virtual screening for 5HT_2A_R utilizes experimentally determined structures including 6A93, 6A94 [38], 6WGT, 6WH4, 6WHA [39], 7RAN [40], 7VOD, 7VOE [41], 7WC4, 7WC5, 7WC6, 7WC7, 7WC8, 7WC9 [42]. As for NET, structures used for virtual screening include 8HFE, 8HFF, 8HFG, 8HFI, 8HFL, 8I3V [43], where 8HFE is modified to ligand-bound complex structures using human serotonin transporter structures as templates [44, 45]. For 5HT_2A_R, the top 2% molecules are extracted, and for NET, the top 1% molecules are extracted. Then, simple drug-likeness filters are applied, with a molecular weight threshold of 550 and a QED [46] threshold of 0.5. The novelty filter excludes molecules that have large ECFP4 similarities to known actives. Known actives are obtained from the ChEMBL database [30], and defined as molecules with a pChEMBEL value > 5, or comments like “active”. The ECFP4 similarity thresholds are set to 0.45 and 0.35 for 5HT_2A_R and NET, respectively.

There is no available experimental structure and active molecules for the HETC domain of TRIP12. The GenPack-generated pockets are used for DrugCLIP virtual screening, and they are downloaded from our website (pocket 1, https://drug-the-whole-genome.yanyanlan.com/drug/Q14669). An updated version of ChemDiv chemical collections was prefiltered with a similar set of rules as **Table S15**. No additional property and novelty filter is applied outside the standard procedure.

All molecules used in these experiments are from chemical collections of ChemDiv, Inc. (https://www.chemdiv.com/), and chemicals are purchased from the TopScience (Tao Shu) Company.

### Functional assays of 5HT_2A_R

The primary screening was conducted via calcium flux assays. All molecules were dissolved in DMSO at 10mM, including the positive control IHCH-7079 [42] and the negative control Risperidone. Calcium flux assays in the agonist mode were conducted by Pharmaron, Beijing, China.

Briefly, Flp-In-CHO-5HT2A cells used in the experiment were cultured in complete medium composed of Ham’s F-12K (Hyclone, SH30526.01), 10% FBS (Gibco, 10999141), Penicillin-Streptomycin (Gibco, 15140122), and Hygromycin B (Invivogen, ant-hg-5) at a final concentration of 600 μg/mL. The cells were maintained under standard conditions at 37°C with 5% CO_2_ to ensure optimal cell density. On the first day of the experiment, the cultured cells were centrifuged and resuspended in an antibiotic-free medium consisting of Ham’s F-12K (Hyclone, SH30526.01) and 10% DFBS (ThermoFisher Scientific, 30067334). Approximately 7,000 cells per well were then seeded into 384-well plates (Corning, 3764) and incubated overnight. The following day, the medium in the 384-well plates was removed, and the cells were thoroughly washed with an assay buffer composed of Hank’s Balanced Salt Solution (HBSS) (Gibco, 14025076) supplemented with 20 mM HEPES (Gibco, 15630080). After washing, 20 μL of assay buffer was left in each well. The 20x Component A from the FLIPR Calcium 6 Assay Kit (Molecular Devices, R8191) was diluted to 2x, and 5 mM probenecid was added. A 20 μL aliquot of this dilution was then added to each well, and the plate was incubated at 37°C for 2 hours. Subsequently, 5x concentrated test solutions of the compounds of interest and a serotonin reference solution were prepared. Using the FLIPR Tetra (Molecular Devices) system, 10 μL of each test compound solution was transferred to the respective wells of the 384-well plate, and the assay results were recorded. Calcium flux assays were repeated three times and recorded relative values were averaged.

Primary hits were defined as molecules that induced > 10% response of the 5-HT reference. These molecules were then verified with radio-ligand comparative binding assays, which were conducted by WuXi Biology. First, 5HT2A-HEK293 cells were cultured, and the cell membranes were harvested to serve as the source of 5HT_2A_R protein, hereafter referred to as the membrane solution, at a concentration of 2.55 mg/mL. According to the experimental design, the test compounds and the reference compound, ketanserin (Sigma-S006), were diluted and 1 μL of each was added to the respective reaction wells. Following this, 100 μL of the membrane solution was added to each well. Next, 100 μL of ^3^H-ketanserin was added to each well to achieve a final concentration of 1 nM. The plates were then sealed and incubated on a shaker at 300 rpm for 1 hour at room temperature. After incubation, 50 μL of 0.3% PEI (Sigma, P3143) solution was added to the Unifilter-96 GF/B filter plates (Perkin Elmer) and incubated for 30 minutes at room temperature. The reaction mixture from each well was then transferred to the filter plates, followed by filtration using a Perkin Elmer Filtermate Harvester. The wells were washed four times with 50 mM Tris-HCl buffer. Subsequently, the filter plates were dried at 50°C for 1 hour. Once dried, the filter plates were sealed at the bottom using Unifilter-96 backing tape (Perkin Elmer), and 50 μL of Microscint 20 cocktail (Perkin Elmer, 6013329) was added to each well. Finally, the top of the plates was sealed with TopSeal-A film (Perkin Elmer). The prepared plates were then placed in a MicroBeta2 Reader (Perkin Elmer) for counting. Radio-ligand comparative binding assays were replicated twice.

Molecules that showed adequate affinities to 5HT_2A_R were further tested with NanoBit assays measuring the recruitment of the β-arrestin2 protein. NanoBit assays were also conducted by Wuxi Biology. On the first day of the experiment, cultured 5HT2A-HEK293 cells were collected. The HEK293 cells were first washed with DPBS solution and then treated with an appropriate amount of 0.25% trypsin-EDTA solution for 5 minutes at 37°C. After digestion, the reaction was quenched by adding an appropriate amount of complete medium, and the mixture was gently mixed. The cells were then centrifuged at 1000 rpm at room temperature to collect the cell pellet. The cells were resuspended to a concentration of 750,000 cells/mL. A 40 μL aliquot of the cell suspension was added to each well of a 384-well plate (Greiner, 781090) and incubated overnight. On the following day, 5 μL of appropriately diluted test samples and control samples were added to each well, followed by the addition of diluted NanoBit assay solution (Promega, N2012). The reaction mixture was incubated at 37°C for 30 minutes. After incubation, the experimental data were read using the Envision2104 (PerkinElmer, 2814243) system. NanoBit assays were replicated twice.

All IC_50_ and K_*d*_ values were fitted with GraphPad Prism.

For structural analysis of hit molecules, molecules are docked to 7WC8 [42] with Glide-SP, and a template of OLC is used for V008-4481 with a RMSD tolerance of 5 Å

### Functional assays of NET

Cells used for NET functional assays included *Escherichia coli* and HEK293F. The *Escherichia coli* strain DH5α was cultured in LB medium (Sigma) at 37 ℃ to generate and amplify plasmids for NET. Mammalian HEK293F cells were maintained in SMM 293-TII medium (Sino Biological) at 37°C with 5% CO2 for protein expression.

The full-length human wild-type NET cDNA (UniProt ID: P23975) was inserted into the pCAG vector using the KpnI and XhoI restriction sites, with an N-terminal FLAG tag. NET overexpression was achieved in HEK293F cells. For transfection, 2 mg of plasmid DNA and 4 mg of polyethylenimine (Polysciences) were pre-incubated in 50 ml of fresh SMM 293-TII medium for 15 minutes before being added to one liter of HEK293F cells at a density of 2.0 × 10^6^ cells/ml. After 48 hours of shaking at 37°C, 5% CO2, and 220 rpm, the cells were collected via centrifugation, resuspended in lysis buffer (20 mM Tris-HCl pH 8.0, 150 mM NaCl), frozen in liquid nitrogen, and stored at -80°C for later use.

For protein purification, the thawed cell pellet was solubilized in lysis buffer containing protease inhibitors (5 μg/ml aprotinin, 1 μg/ml pepstatin, 5 μg/ml leupeptin; Amresco) and 2% (w/v) DDM (Anatrace) at 4°C for 2 hours, followed by centrifugation at 20,000 g at 4°C for 1 hour. The resulting supernatant was applied to anti-FLAG M2 resin (Sigma), which was washed with 15 column volumes (CV) of buffer (20 mM Tris-HCl pH 8.0, 150 mM NaCl, 0.02% (w/v) DDM). The protein was eluted with 6 CV of the wash buffer containing 0.4 mg/ml FLAG peptide at 4°C. The eluted protein was concentrated and further purified by size-exclusion chromatography using a Superose 6 Increase 10/300 GL column (GE Healthcare) in buffer (20 mM Tris-HCl pH 8.0, 150 mM NaCl, 0.02% (w/v) DDM). The peak fractions were collected and concentrated for subsequent experiments.

Then, purified NET protein was reconstructed into liposomes to form proteoliposomes. The *E. coli* polar lipid extract (Avanti), with 20% (wt %) cholesterol added, was resuspended to 20 mg/ml in buffer A (25 mM HEPES pH 7.4, 150 mM KCl). This mixture underwent ten freeze-thaw cycles using liquid nitrogen and was then extruded 21 times through 0.4 μm polycarbonate membranes (GE Healthcare). The resulting liposomes were pre-treated with 1% n-octyl-β-D-glucoside (β-OG; Anatrace) for 30 minutes at 4°C. They were then incubated with 200 μg/ml of purified NET protein (wild-type or mutants) for 1 hour at 4°C. To remove the detergents, the mixture was treated overnight with 250 mg/ml Bio-Beads SM2 (Bio-Rad) at 4°C, followed by an additional 1-hour incubation with 100 mg/ml Bio-Beads SM2. After five more freeze-thaw cycles and 21 additional extrusion passes, the proteoliposomes were collected by ultracentrifugation at 100,000 g for 1 hour at 4°C, washed twice, and resuspended to 100 mg/ml in buffer A for the subsequent uptake assay.

Each uptake assay was conducted by adding 2 μl of proteoliposomes to 96.5 μl of buffer B (25 mM HEPES pH 7.4, 150 mM NaCl) along with 0.5 μl (0.5 μCi, 12.3 Ci/mmol) of Levo-[7-3H]-Norepinephrine and 1 μl of 50 μM valinomycin. To assess the single-point inhibitory activity of the screened small molecules, proteoliposomes were incubated with these compounds, while Desipramine and Bupropion were used as positive controls for NET inhibition. All inhibitors were added at a concentration of 1 μM in a volume of 1 μl. The uptake of the radiolabeled substrates was halted after 60 seconds by rapidly filtering the solution through 0.22 μm GSTF filters (Millipore) and washing with 2.5 ml of ice-cold buffer B. Filters were then incubated with 0.5 ml of Optiphase HISAFE 3 (PerkinElmer) overnight, and radioactivity was measured using a MicroBeta2® Microplate Counter (PerkinElmer). For IC_50_ determination of antidepressants, proteoliposomes were pre-incubated with varying concentrations of the drugs for 30 minutes before the addition of isotope-labeled substrates. IC_50_ values were calculated using GraphPad Prism 8, applying non-linear regression to fit the data to the equation:

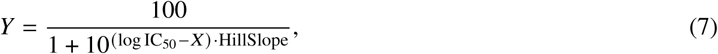

with option: ‘log(inhibitor) vs. normalized response—Variable slope’. X represents the log of the inhibitor concentration, Y represents the normalized response (ranging from 100% to 0%), and HillSlope starts with an initial value of -1.

All experiments were conducted in triplicate using biologically independent samples. Data were normalized to the wild-type protein to express values relative to 100%. Non-specific binding was accounted for by using control liposomes without protein insertion, ensuring that only specific interactions were measured.

### Synthesis of 0086-0043 and Y510-9709

Both molecules were synthesized by Bellen Chemistry Company.

For Y510-9709 (**5-(4-chlorophenyl)-2,3-dihydrothiazolo[2,3-b]thiazol-4-ium bromide**), first synthesize compound 2 (**1-(4-chlorophenyl)-2-((4,5-dihydrothiazol-2-yl)thio)ethan-1-one**). To a solution of compound 1 (**2-bromo-1-(4-chlorophenyl)ethan-1-one**) (10.0 g, 42.8 mol, 1.0 eq) and **thiazolidine-2-thione** (5.1 g, 42.8 mmol, 1.0 eq) in EtOH (150 mL) and DMF (50 mL) was added TEA (4.3 g, 42.8 mol, 1.0 eq). The reaction mixture was stirred at room temperature for 2 h. HPLC showed no compound 1 remained. The reaction mixture was poured into crushed ice and filtered to give compound 2 (9.6 g, 82.5%) as a yellow solid. 1H NMR (300 MHz, CDCl3): δ ppm 8.00 – 7.90 (m, 2H), 7.50 – 7.40 (m, 2H), 4.62 (s, 2H), 4.17 (t, J = 8.1 Hz, 2H), 3.43 (t, J = 7.8 Hz, 2H). LCMS: 272.0 ([M+H]+).

Then, The solution of compound 2 (2.5 g, 9.2 mmol, 1.0 eq) in 30% HBr in AcOH (25 mL) was stirred at 120 °C for 3 h. TLC and HPLC showed no compound2 remained. The reaction was allowed to be cooled to room temperature and concentrated in vacuo to give the residue, which was triturated with MeOH (7.5 mL) and filtered to give Y510-9709 (1.1 g, 35.7%) as an off-white solid. 1H NMR (400 MHz, DMSO-d6): δ ppm 7.90 (s, 1H), 7.68 (s, 4H), 4.70 (t, J = 8.0 Hz, 2H), 4.10 (t, J = 8.4 Hz, 2H). LCMS: 254.0 ([M-Br]+).

For 0086-0043(**2-(2-oxo-2-phenylethyl)isoquinolin-2-ium chloride**), The solution of **2-chloro-1-phenylethan-1-one** (2.0 g, 12.9 mol, 1.0 eq) and **isoquinoline** (1.7 g, 12.9 mmol, 1.0 eq) in ACN (12 mL) was stirred at room temperature for 16 h. HPLC showed no **2-chloro-1-phenylethan-1-one** remained. The reaction mixture was filtered to give 0086-0043 (1.3 g, 35.4%) as an off-white solid. 1H NMR (400 MHz, DMSO-d6): δ ppm 10.06 (s, 1H), 8.76 (d, J = 6.8 Hz, 1H), 8.69 (d, J = 6.8 Hz, 1H), 8.56 (d, J = 8.4 Hz, 1H), 8.43 (d, J = 8.4 Hz, 1H), 8.39 – 8.28 (m, 1H), 8.20 – 8.04 (m, 3H), 7.81 (t, J = 7.6 Hz, 1H), 7.69 (t, J = 7.6 Hz, 2H), 6.66 (s, 2H). LCMS: 248.1 ([M-Cl]+).

### The structure determination of NET and its inhibitors

For cryo-EM samples, 4μl purified NET protein was applied to glow-discharged Quantifoil holey carbon grids (Quantifoil Au R1.2/1.3, 300 mesh). Protein was concentrated to approximately 10 mg/ml and separately incubated with 2 mM Y510-9709 or 0086-0043 for 30 min before freezing. After applying the protein, the grids were blotted for 3 s with 100% humidity at 4 °C and plunge frozen in liquid ethane cooled by liquid nitrogen with Vitrobot (Mark IV, Thermo Fisher Scientific).

Cryo-EM data were collected on a 300 kV Titan Krios G3i equipped with a Gatan K3 detector and a GIF Quantum energy filter (slit width 20 eV). The defocus values ranged from -1.5 to -2.0 μm. Each stack of 32 frames was exposed for 2.56 s, and the exposure time of each frame was 0.08 s. The micrographs were automatically collected with AutoEMation program [47] in super-resolution counting mode with a binned pixel size of 1.083 Å. The total dose of each stack was about 50 e^−^/Å^2^. All 32 frames in each stack were aligned and summed using the whole-image motion correction program MotionCor2 [48].

All dose-weighted micrographs were manually inspected and imported into cryoSPARC [49]. Micrographs with an estimated CTF resolution worse than 4 Å were excluded during exposure curation. CTF parameters were estimated using patch-CTF. They were used for initial good templates generation via 2D classification. Initial good templates were generated via 2D classification, using the previously reported NET structure [50] (NET–DSP, PDB code: 8FHI) as a reference. The Template Picker tool was used for all particle picking tasks. For the NET_Y510-9709 and NET_0086-0043 datasets, 3,204,486 and 9,008,886 particles were extracted from 2,918 and 4,687 micrographs, respectively. Particles were initially extracted with a box size of 192 and then cropped to 128 to speed up calculations. The initial good reference for 3D classification was derived from the NET-DSP dataset, while bad references were generated using the graphical user interface (GUI) of UCSF ChimeraX [51]. Global pose estimation was performed using Non-uniform refinement, followed by local refinement for the first round of local pose assignment. A second round of local pose estimation was conducted using 3D classification (without image alignment), followed by another round of local refinement (**Fig**.**S8**). This process yielded 507,444 and 506,286 particles representing the inward-open conformation, resulting in resolutions of 2.87 Å for NET_Y510-9709 and 2.98 Å for NET_0086-0043, respectively. The atomic coordinates of NET in the presence of Y510-9709 or 0086-0043 have been deposited in the Protein Data Bank (http://www.rcsb.org) under accession codes 9JEL and 9JF3. The corresponding electron microscopy maps are available in the Electron Microscopy Data Bank (https://www.ebi.ac.uk/pdbe/emdb/) under accession codes EMD-61420 and EMD-61426.

### The training and inference of the GenPack generative model

We have developed a GenPack model that operates within a continuous parameter space, incorporating a noise-reduced sampling strategy inspired by MOLCRAFT [52]. Unlike full-atom approaches, our method focuses solely on the given backbone atoms to minimize the impact of potential structural variations between *apo* and *holo* states of the proteins. We meticulously curate a dataset comprising 14,616 protein-ligand pairs from the PDBbind database, which we divide into a training set of 13,137 pairs and a validation set of 1,479 pairs (Supplementary Materials 3). Additionally, we use 101 protein-ligand pairs from the DUD-E database as our test set. To prevent data leakage, we excluded all proteins from the training and validation sets that share a FLAPP similarity score greater than 0.9 with any target in the test set. FLAPP [53] is a tool used to estimate the structural similarity (alignment rate) between two pockets. Pockets are defined by extracting backbone atoms within a 10 Å radius of the ligands. The training is conducted on a single NVIDIA A100 GPU with a learning rate of 5e-4 for 60 epochs, resulting in our pocket location optimization model.

During inference, Fpocket [54] is initially employed to detect pockets approximately 10 Å in size, after which our SBDD model generates potential ligand molecules conditioned on backbone atoms only. Subsequently, side-chain atoms are introduced to the complex structure, and the complex structures are relaxed with Prime software in the Schrodinger Suite. The protein residues with at least one heavy atom within a 6 Å radius of the generated ligands are selected as the final pocket region. This approach ensures a focus on critical interactions within the binding site while reducing noise and irrelevant structures, thereby facilitating accurate pocket detection.

### Evaluating the effectiveness of GenPack model

To evaluate the effectiveness of the GenPack model, we conducted experiment on the targets of DUD-E.

We conducted two types of experiments to evaluate the effectiveness of the GenPack algorithm in refining protein structures.

In the first experiment, we utilized AlphaFold-predicted structures of protein targets, optimized using GenPack, to perform virtual screening against the DUD-E dataset. The screening performance was assessed using the Enrichment Factor (EF) metric. We identified AlphaFold2 (AF2) structures corresponding to the UniProt entries of DUD-E targets in the AlphaFold database, yielding a total of 96 targets. For the GenPack results, five conformations were sampled for each target, and the best-performing conformation was selected for evaluation. The detailed results are provided in **Table S11**.

Additionally, we evaluated the performance of GenPack on *apo* structures. The corresponding results are also presented in **Table S10**. The *apo* structures were obtained from a previous research [22] and encompass 27 protein targets included in the DUD-E dataset.

In the second experiment, we assessed the structural accuracy of GenPack-refined proteins through redocking. Specifically, we docked the original ligand back into the GenPack-generated protein structure and measured the Root-Mean-Square Deviation (RMSD) between the redocked and the original ligand conformations. Results presented in **Table S12** and **S13**. For pockets without GenPack optimization, five docking poses were generated, and the best one was selected. For GenPack-optimized pockets, five pocket conformations were generated; for each conformation, only a single docking pose was used. The best result among these five pocket conformations was then selected.

We also measure the correlation of the pockets localization, sidechain accuracy and docking or virtual screening effects, shown in **Fig. S10**. We show in **Fig. S10A** the impact of the GenPack method on pocket localization performance, measured by Intersection over Union (IoU), and on virtual screening effectiveness compared to *holo* structure. Pocket localization ability is assessed by the IoU between the predicted pocket and the corresponding *holo* pocket. Here, the virtual screening metric EF1% represents the reduction in enrichment factor when using Fpocket prediction of AlphaFold structures relative to *holo* structures. As the IoU with the *holo* structure increases, the reduction in EF1% correspondingly decreases. The GenPack method enables Fpocket results more spatially aligned with the *holo* pockets, thereby narrowing the performance gap in EF1%.

**Fig. S10B** illustrates the relationship between side-chain RMSD of the predicted pocket and the reduction in EF1%. The observed p-value is relatively large, suggesting that the correlation is not statistically significant within the DUD-E dataset. Moreover, the GenPack method does not substantially alter the distribution of side-chain RMSD between Fpocket-predicted pockets and their corresponding *holo* pockets.

**Fig. S10C** and **D** examine the relationship between structural pocket accuracy and Glide-SP docking performance, as measured by ligand RMSD. In **Fig. S10C**, the correlation between pocket IoU (with respect to *holo* pockets) and docking accuracy is evaluated, with both docking grid centers and pocket definitions obtained through structural alignment. The results suggest no significant difference in ligand docking pose RMSD as a function of pocket localization accuracy. Similarly, **Fig. S10D** investigates the impact of side-chain RMSD of the predicted pocket (relative to the *holo* structure) on docking accuracy. The analysis reveals no evident correlation between ligand RMSD and variations in side-chain conformations, indicating that deviations in side-chain positioning have minimal effect on docking pose accuracy.

### Protein expression and purification of TRIP12

The plasmid encoding human TRIP12 (442–1992) gene was cloned into the pGEX-4T-1 vector, which was fused with an N-terminal GST tag followed by an HRV 3C protease cleavage site. This construct was synthesized and optimized for Escherichia coli overexpression by GenScript (Nanjing, China).

The recombinant plasmid was transformed into BL21 (DE3) cells and then cultured in Luria Broth media containing 50 µg/mL ampicillin at 37°C. When the optical density of the culture reached 0.6–0.8, protein expression was induced by adding 0.4 mM IPTG at 16°C. After overnight incubation, cells were harvested by centrifugation at 5000 × *g* for 30 min at 4°C and resuspended in the lysis buffer (50 mM HEPES, 150 mM NaCl, pH 7.5). Cells were then lysed by ultrasonication and the lysate was centrifuged at 12500 × *g* for 30 min at 4°C to remove precipitates.

The supernatant was applied to Glutathione beads for 2 h at 4°C, and target proteins fused with GST tag were eluted with elution buffer (50 mM HEPES, 150 mM NaCl, 30 mM Glutathione, pH 7.5). After removing the GST tag with HRV 3C protease, proteins were further purified with ion exchange chromatography (HiTrap Heparin column, GE Healthcare) followed by size exclusion chromatography (Superdex 6 Increase column, GE Healthcare).

### Surface Plasmon Resonance (SPR) analysis

Surface plasmon resonance experiments were performed using a Biacore 8k (Cytiva) at 25°C. TRIP12 was immobilized on a CM7 sensor chip (Cytiva) using standard amine coupling chemistry. Briefly, the carboxymethylated dextran surface was activated with a 1:1 mixture of 0.4 M EDC (1-ethyl-3-(3-dimethylaminopropyl)carbodiimide) and 0.1 M NHS (N-hydroxysuccinimide) for 420 s. The protein (50 µg/mL in 10 mM sodium acetate, pH 4.0) was then injected over the activated surface until reaching approximately 12000 response units (RU). Remaining activated groups were blocked with 1 M ethanolamine-HCl (pH 8.5). A reference flow cell was prepared by activating and blocking the surface without protein immobilization.

Compounds were dissolved in DMSO and diluted in running buffer (PBS pH 7.4, containing 0.05% Tween-20 and 2% DMSO) to maintain a constant DMSO concentration. To account for bulk refractive index changes caused by DMSO, solvent correction was performed using a series of running buffer containing four DMSO concentrations ranging from 0.5% to 4%. Concentration ranges were adjusted for each compound to enable accurate determination of *K*_d_ values. Different compounds required different concentration series depending on their binding characteristics. A serial dilution series of each compound was injected over the immobilized protein and reference surfaces at a flow rate of 30 µL/min.

In the screening experiments, single-cycle kinetics was employed with a series of increasing compound concentrations injected sequentially with a contact time of 120 s followed by a 240 s dissociation phase after the final injection. For affinity validation experiments, multi-cycle kinetics was performed where each compound concentration was injected individually with a contact time of 120 s and a dissociation time of 200 s before regeneration of the sensor surface. After solvent correction was performed, sensorgrams were referenced by subtracting both reference flow cell and blank buffer injection responses. For both single-cycle and multi-cycle kinetic experiments, steady-state binding responses were fitted to a 1:1 binding model using Biacore Evaluation Software to determine the equilibrium dissociation constant (*K*_d_).

### Determine the enzyme activity of TRIP12 with the *in vitro* ubiquitination assay

*In vitro* ubiquitination assays were performed with a specific K48diUb^prox-K29^ substrate, as previously described [55]. In brief, 0.5 µM Uba1, 4 µM Ubch7, 0.25 µM TRIP12, 2 µM fluorescent K48-linked diUb with lysine to arginine mutation at the distal LYS29 site and keeping the proximal LYS29 unchanged (named K48diUb^prox-K29^), 80 µM WT Ub, and either varying concentrations of E599-0223 or G935-3912 (dissolved in DMSO) or DMSO alone (as control) were mixed at 37°C for 2 minutes in the reaction buffer (50 mM HEPES, pH 7.5, 150 mM NaCl, 10 mM MgCl_2_, and 5 mM ATP). The reaction was terminated with 4× SDS sample buffer with DTT, and analyzed by SDS-PAGE followed by fluorescence imaging and Coomassie Brilliant Blue dye (Bio-Rad).

### E1∼Ub and E2∼Ub thioester formation assay with fluorescent Ub

The conditions for the E1∼Ub thioester formation assay are as follows: 0.5 µM Uba1, 10 µM fluorescent Ub, and either 400 µM E599-0223 or G935-3912 dissolved in DMSO (or DMSO alone as control) were mixed at 37°C for 5 minutes in the reaction buffer (50 mM HEPES, pH 7.5, 150 mM NaCl, 10 mM MgCl_2_, and 5 mM ATP). The reaction was terminated with 4× SDS sample buffer, with or without DTT, and analyzed by SDS-PAGE followed by fluorescence imaging and Coomassie Brilliant Blue dye (Bio-Rad). The E2∼Ub thioester formation assay was performed under the same conditions, except that 5 µM Ubch7 was additionally included in the reaction.

### Pocket Detection for all AlphaFold2 predicted human proteins

The AlphaFold DB [56, 57] contains predicted structures for 20,504 human proteins identified by UniProt accessions. Among these, 208 proteins are larger than 2500 amino acids (AAs), and their Pairwise Alignment Error (PAE) cannot be accessed through the official website. Consequently, only 20,296 proteins are used for pocket detection. Not all AlphaFold2 predictions are accurate. Two types of inaccuracies can be avoided by examining the pLDDT and PAE scores. First, we remove all residues with a pLDDT score below 50. The remaining structures exhibit high local accuracy, but the interactions between protein domains may still be incorrect. To address this, the PAE is symmetrized and used as precomputed metrics for agglomerative clustering. The average linkage method is applied, and the PAE threshold for clustering is set at 15 Å. Each cluster is then regarded as a confidently predicted protein super-domain, and protein fragments shorter than 10 AAs are removed to ensure stability during refinement. From the 20,296 proteins, we have identified 24,692 super-domains, covering 17,188 proteins (69.6%).

For each super-domain, we utilize two methods to detect potential pockets. First, we implement a template-based structural alignment approach. Each super-domain is aligned with proteins from the PDBbind database [26, 27]. When a local structure of the super-domain exhibited high structural similarity to a known pocket from PDBbind, it is considered a likely pocket. Specifically, TM-align [58] is used for structural alignment, with a TM-score threshold of 0.6 to ensure significant overall similarity. The corresponding ligands from PDBbind are mapped to the identified pocket location in the super-domain using a rotation matrix, thereby confirming the pocket. We then calculate the local alignment IoU (intersection over union) for the pocket, defined as the ratio of the number of aligned amino acids in the pocket to the number of the union of amino acids in both the super-domain and the PDBbind protein pockets. Alignments with an IoU exceeding 0.6 are retained. Since all super-domains are single-chain proteins, only proteins from PDBbind with single-chain pockets are used for template matching. We also exclude ligand-receptor pairs from the PDBbind database where the ligand contains more than 800 atoms. In addition to the approach above, for each super-domain, Fpocket software [54] is used for pocket detection. However, the accuracy of pocket detection using Fpocket alone is limited, and the side-chain conformation of the *apo* pocket is not suitable for molecular docking. To address this, we adopt the proposed GenPack method to refine the pocket.

### The chemical library for the genome-wide virtual screening

ZINC database is pre-filtered by anodyne reactivity and lead-like properties (molecular weight is no less than 200 and up to 500, logP is up to 5). The resulting subset contains 2,782 tranches, and over 609 million protomers are downloaded from ZINC20 [59]. Enamine REAL database is downloaded from VirtualFlow [60] in the format of PDBQT. The whole database contains 46570 tranches, over 1337 million protomers. Both databases are filtered by cutoff rules for molecular properties calculated from SMILES and structural alert patterns using RDKit. Molecules of properties meeting the rules in **Table S15** are kept for subsequential research. For ZINC, SMILES strings are matched to 3D structures in PDBQT by ZINC id. For REAL, SMILES strings are first extracted from remarks in PDBQT files; if errors like syntax errors due to the letter ’q’ in SMILES occurred, they are then converted from PDBQT structures via Open Babel. A regular expression filter is applied to REAL to exclude PDBQT files with overflowed atom coordinate digits.

### The genome-wide virtual screening

All pockets and molecules are pre-encoded with DrugCLIP models. Then cosine similarities of their embeddings are calculated with Pytorch [61] with 8 A100 GPUs. Then, scores from 6 models and multiple pocket replicas are ensembled as discussed previously. The top 100,000 molecules for each pocket are obtained, and clustered into around 100 clusters with an ECFP4 cut-off of 0.15. Finally, the remaining molecules are docked to the pocket replica with the highest fitness with Glide-SP software from the Schrodinger Suite. Only molecules with a DrugCLIP Zscore > 4 and Glide Score < -6 are included in the final database.

## Supplementary Tables and Figures

**Fig. S1.**
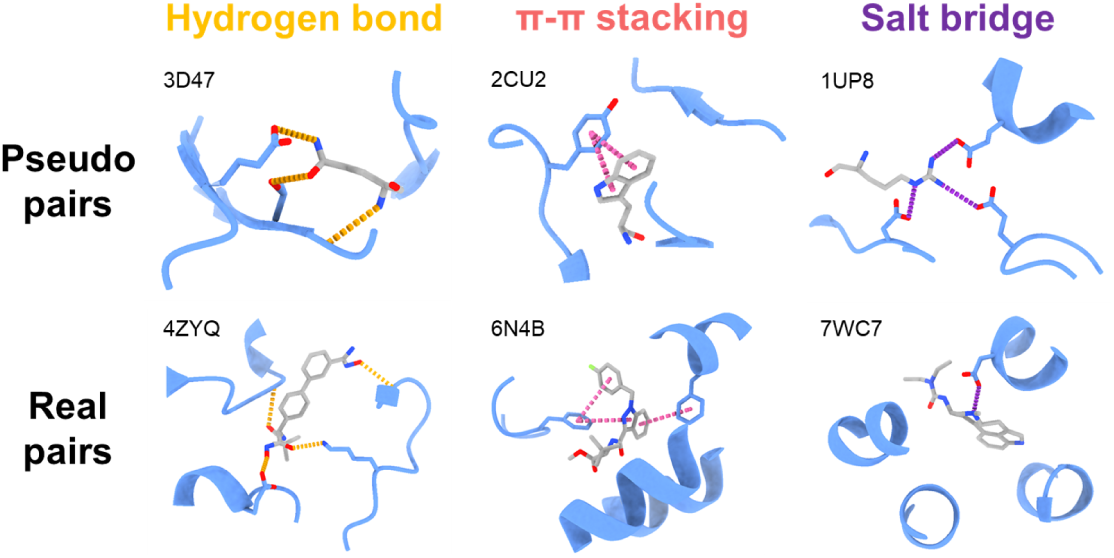
Visualizations of non-covalent interactions shared by both real protein-ligand pairs and pseudo protein-ligand pairs.

**Fig. S2.**
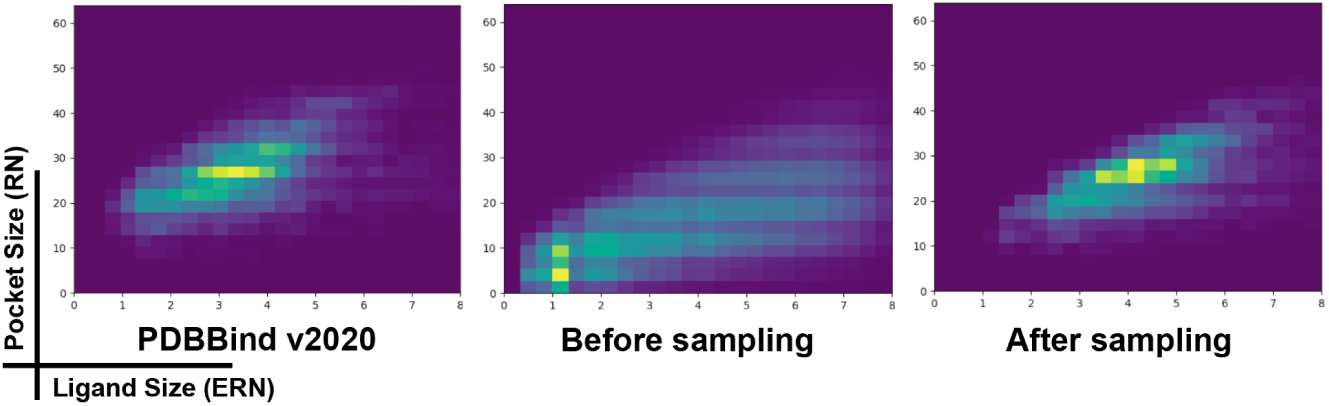
The joint distributions of pocket size and ligand size are examined for the PDBBind dataset, our ProFSA dataset before applying stratified sampling, and the ProFSA dataset after stratified sampling.

**Fig. S3.**
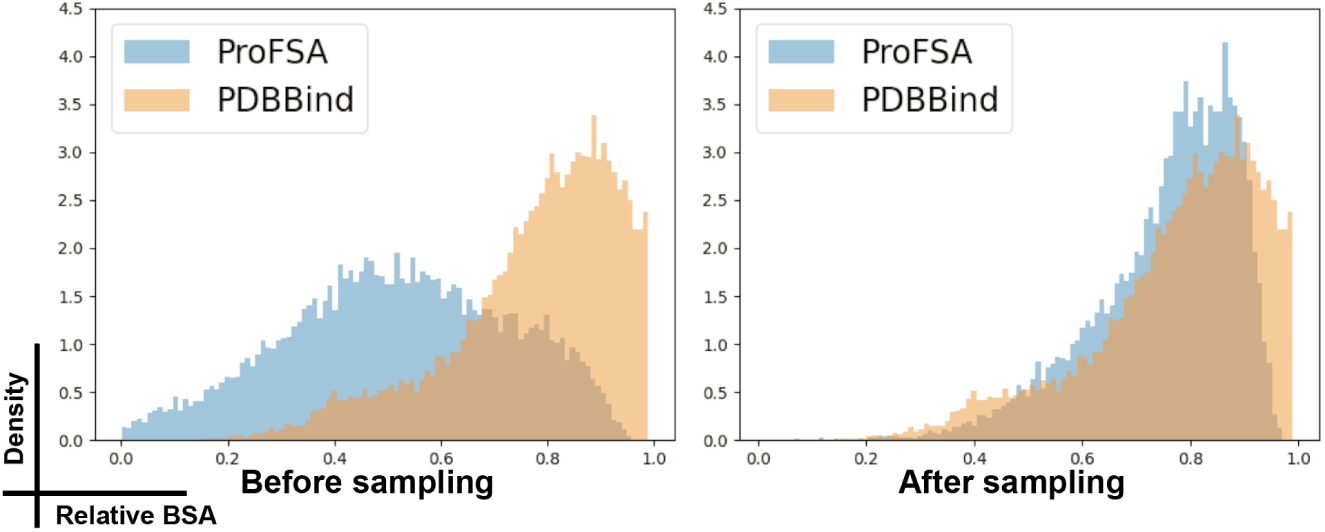
Comparisons between the ProFSA dataset and the PDBBind dataset are made based on the distributions of relative Binding Surface Area (rBSA) for ligand-pocket pairs

**Fig. S4.**
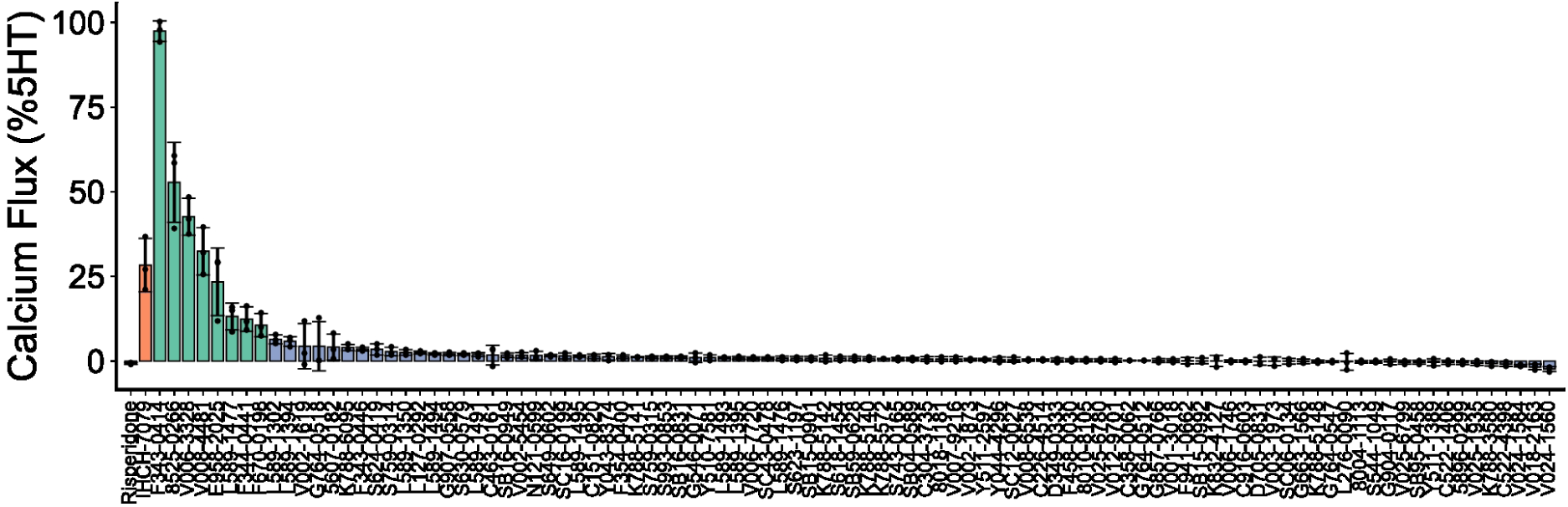
Wet-lab validations of DrugCLIP with 5HT_2A_R. The screening results of 78 DrugCLIP identified molecules using calcium flux assays for 5HT_2A_R agonist at a concentration of 10 µM. Eight molecules showed signals larger than 10%. Orange color indicates positive controls, and green color indicates hit molecules.

**Fig. S5.**
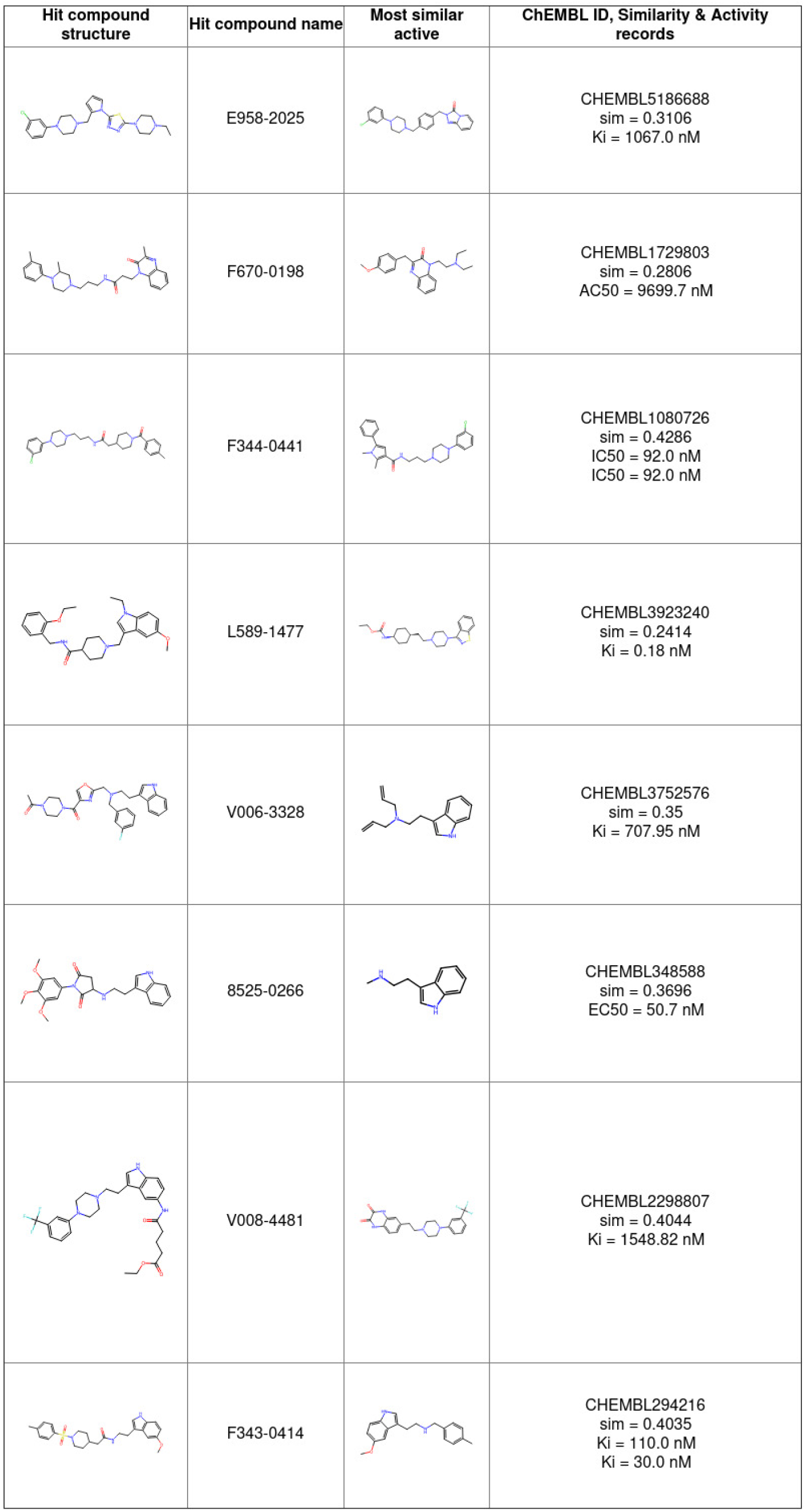
Primary hit molecules of 5HT_2A_R and the known actives with the largest similarity scores. All similarity scores were calculated with Canvas software from the Schrodinger Suite with the ECFP4 fingerprint.

**Fig. S6.**
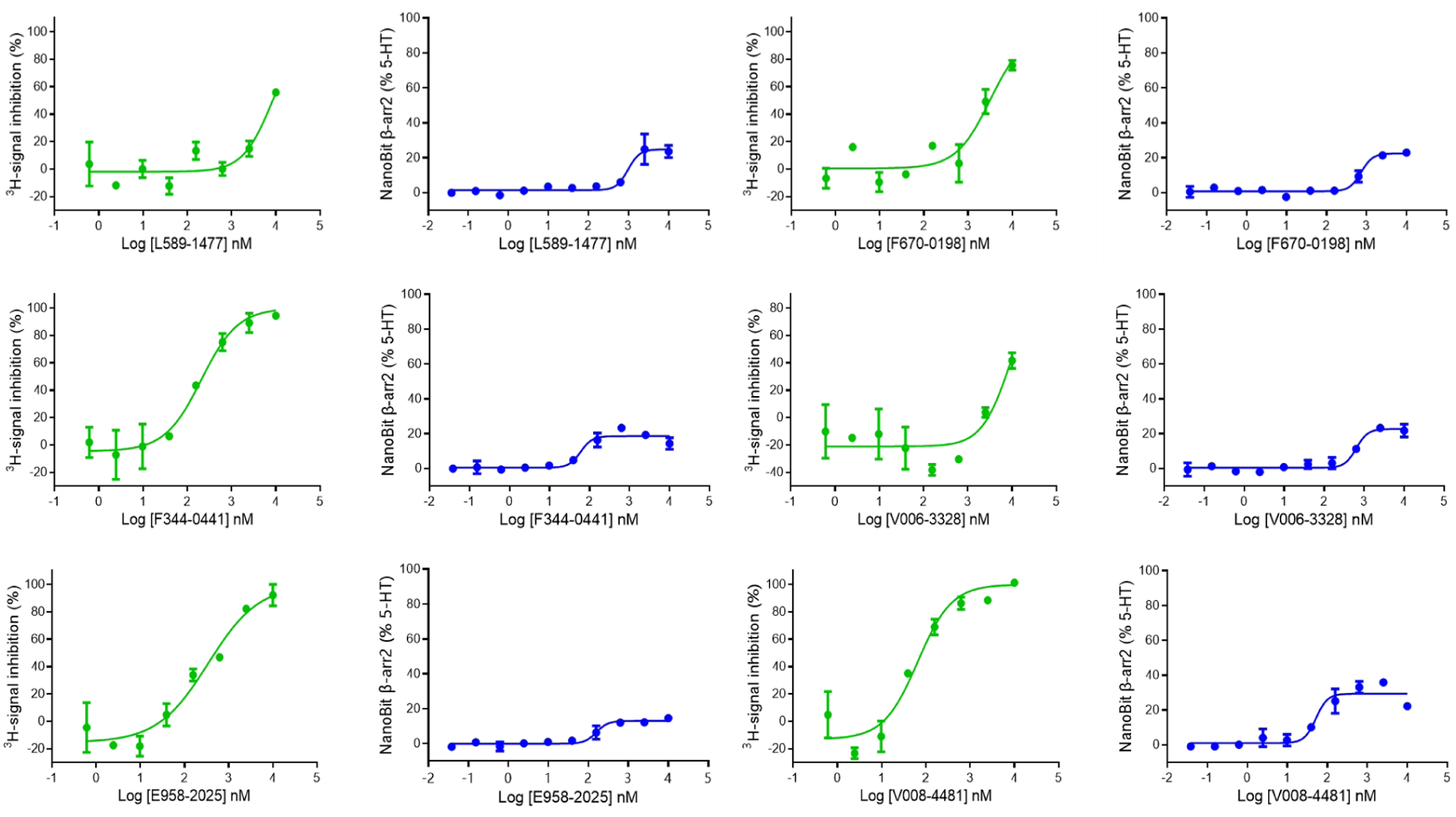
Dosage response curves of primary hits of 5HT_2A_R in radio-ligand competitive binding assays and NanoBit assays.

**Fig. S7.**
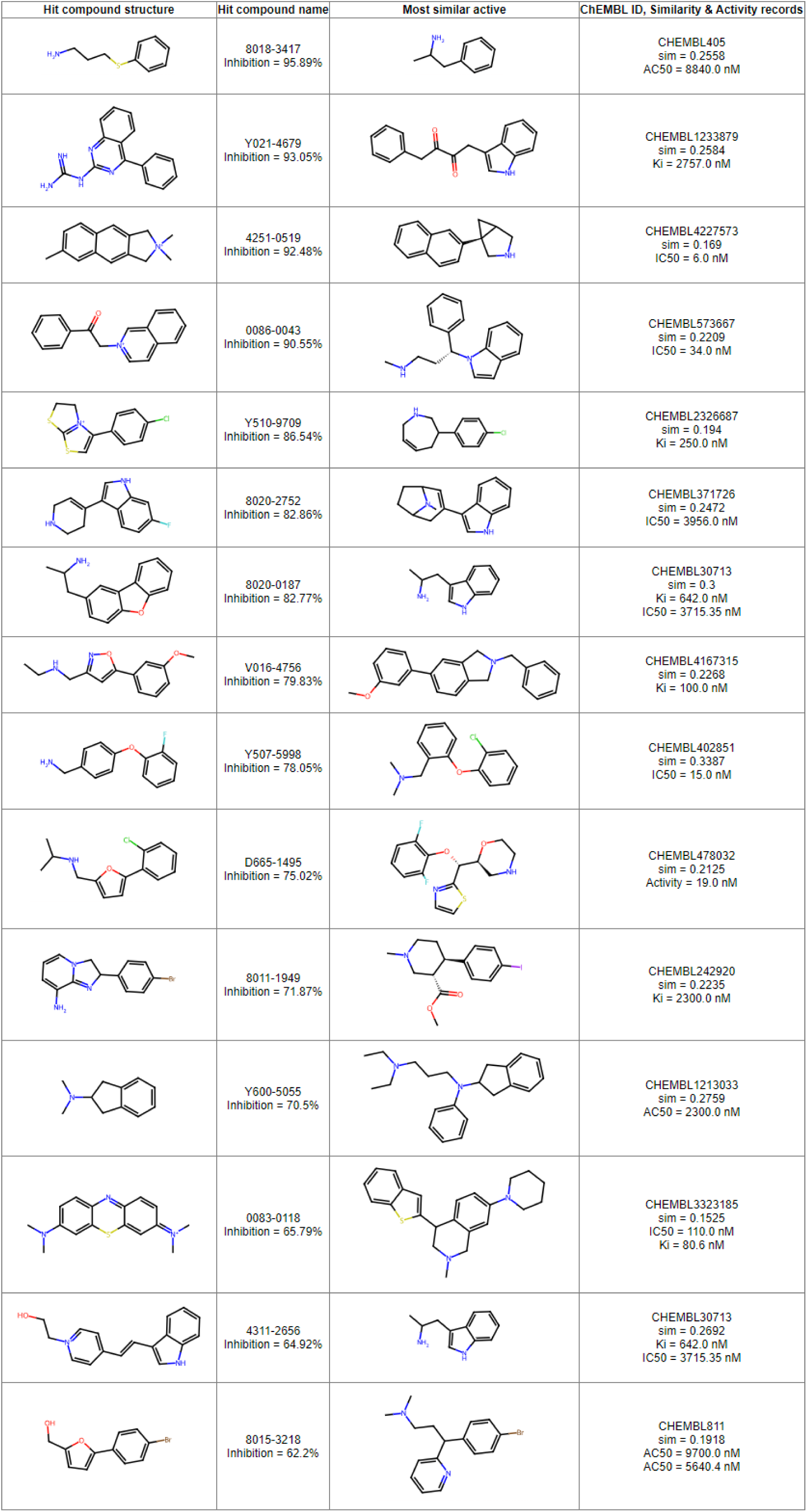
Primary hit molecules of NET and the known actives with the largest similarity scores. All similarity scores were calculated with Canvas software from the Schrodinger Suite with the ECFP4 fingerprint.

**Fig. S8.**
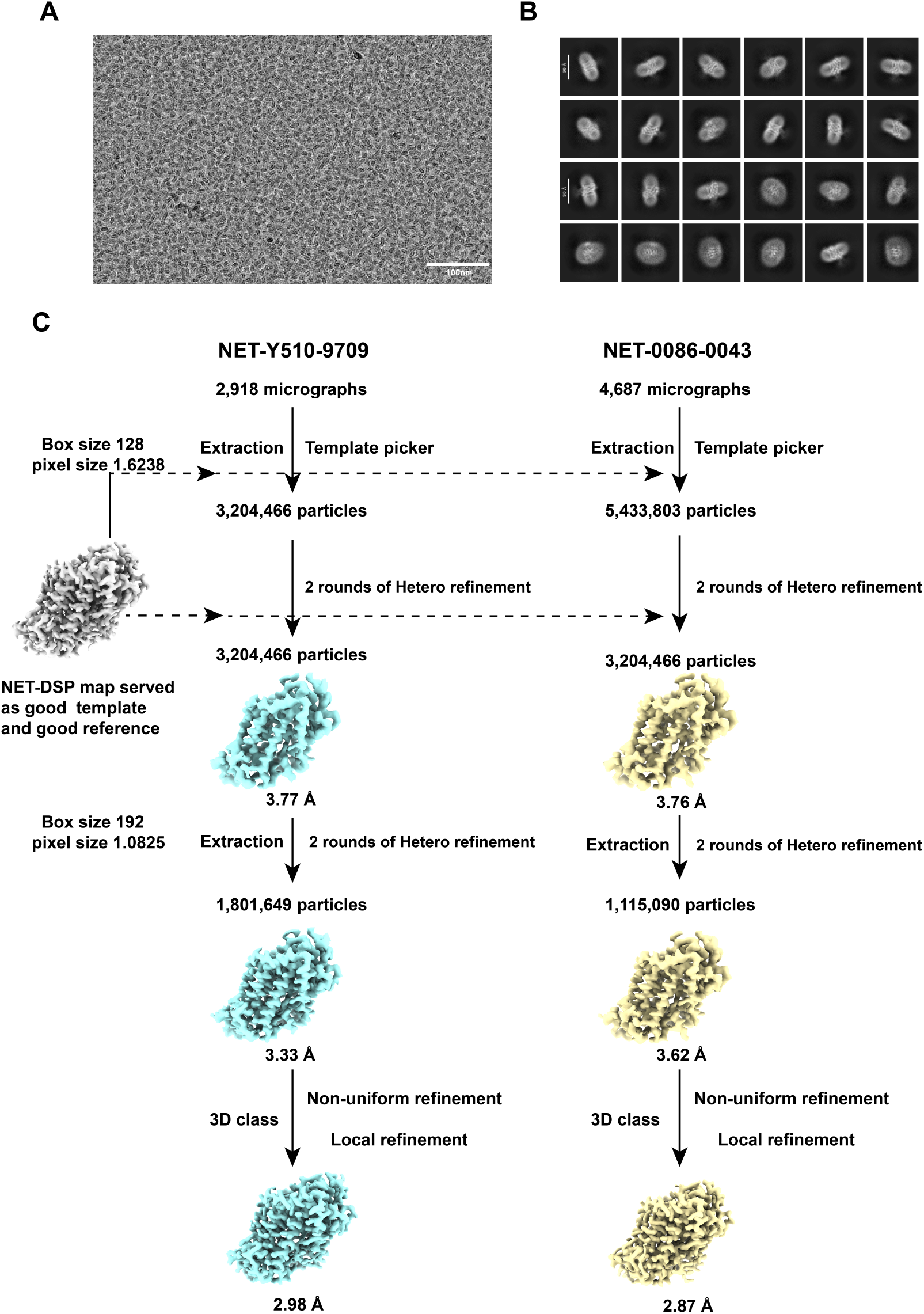
Data processing of NET datasets. (A-B) Representative micrograph and 2D class averages of NET. (C) The flowchart for the data processing of NET bound to Y510-9709 or 0086-0043.

**Fig. S9.**
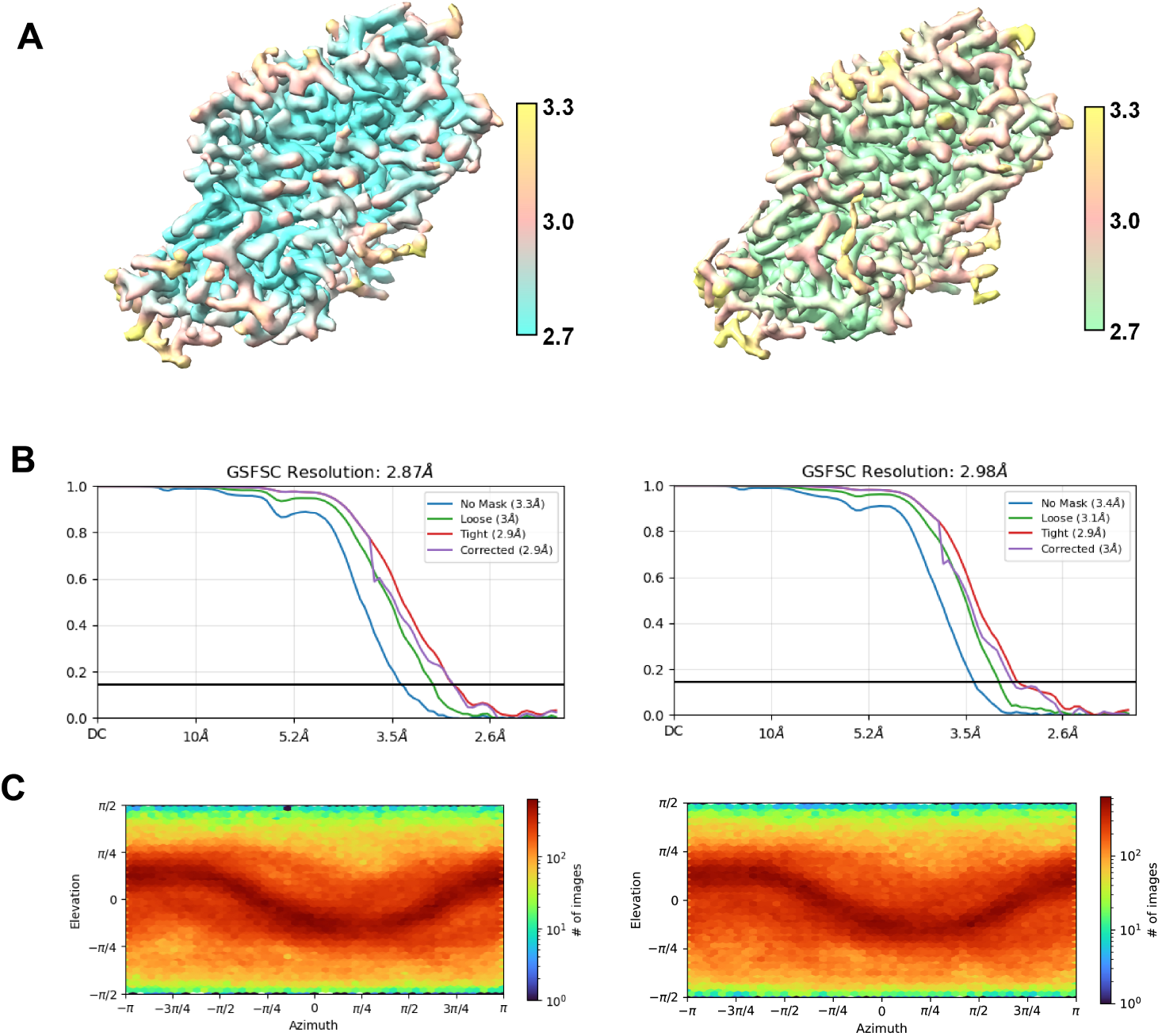
Cryo-EM analysis of NET datasets. Left panel: NET bound to Y510-9709; Right panel: NET bound to 0086-0043. Various assessments of the cryo-EM reconstruction are presented. These include (A) local resolution maps; (B) gold-standard Fourier shell correlation (FSC) curves; (C) angular distribution of the particles used for the final reconstruction.

**Fig. S10.**
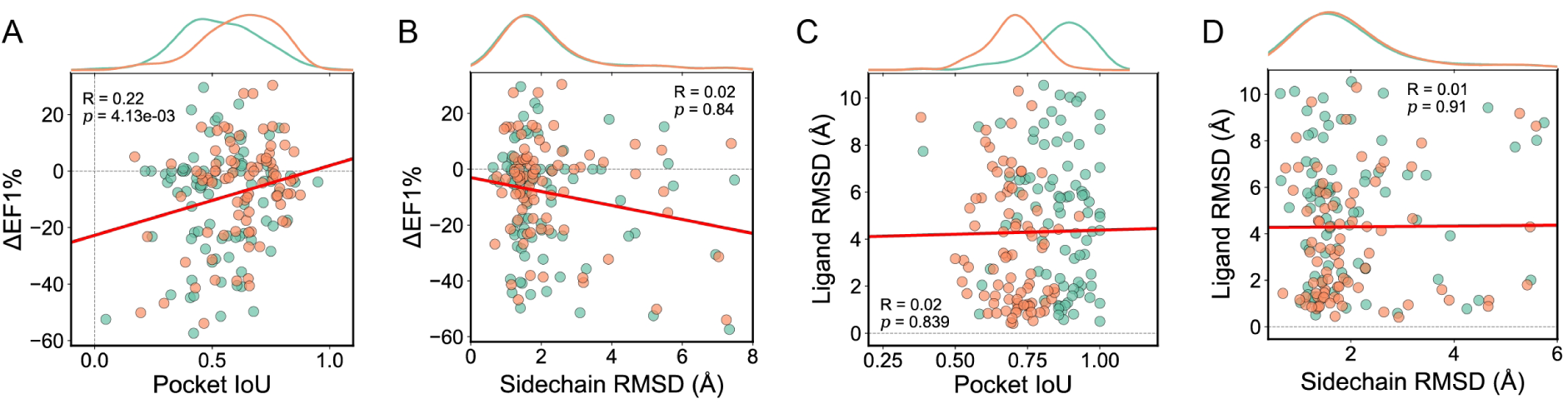
Analysis of the impact of sidechain accuracy and pocket definition on virtual screening and molecular docking performance. (A) Correlation between pocket IoU compared with *holo* pockets to EF1% performance decreases. Green dots indicate samples of Fpocket predictions, while orange dots indicate refined pockets by GenPack. The curves at the top of the plot represent the marginal distribution of pocket IoU. (B) Correlation between pocket sidechain RMSD compared with *holo* pockets to EF1% performance decreases. Green dots indicate samples of Fpocket predictions, while orange dots indicate refined pockets by GenPack. The curves at the top of the plot represent the marginal distribution of sidechain RMSD. (C) Correlation between pocket IoU compared with *holo* pockets to Glide-SP docking accuracy measured by ligand RMSD. Green dots indicate samples using AlphaFold2 predictions as receptors, while orange dots indicate docking with AlphaFold2 structures refined by GenPack. Both docking grid centers and pocket definitions are acquired via structural alignments. The curves at the top of the plot represent the marginal distribution of pocket IoU. (D) Correlation between pocket sidechain RMSD compared with *holo* pockets to Glide-SP docking accuracy measured by ligand RMSD. Green dots indicate samples using AlphaFold2 predictions as receptors, while orange dots indicate docking with AlphaFold2 structures refined by GenPack. Both docking grid centers and pocket definitions are acquired via structural alignments. The curves at the top of the plot represent the marginal distribution of sidechain RMSD.

**Fig. S11.**
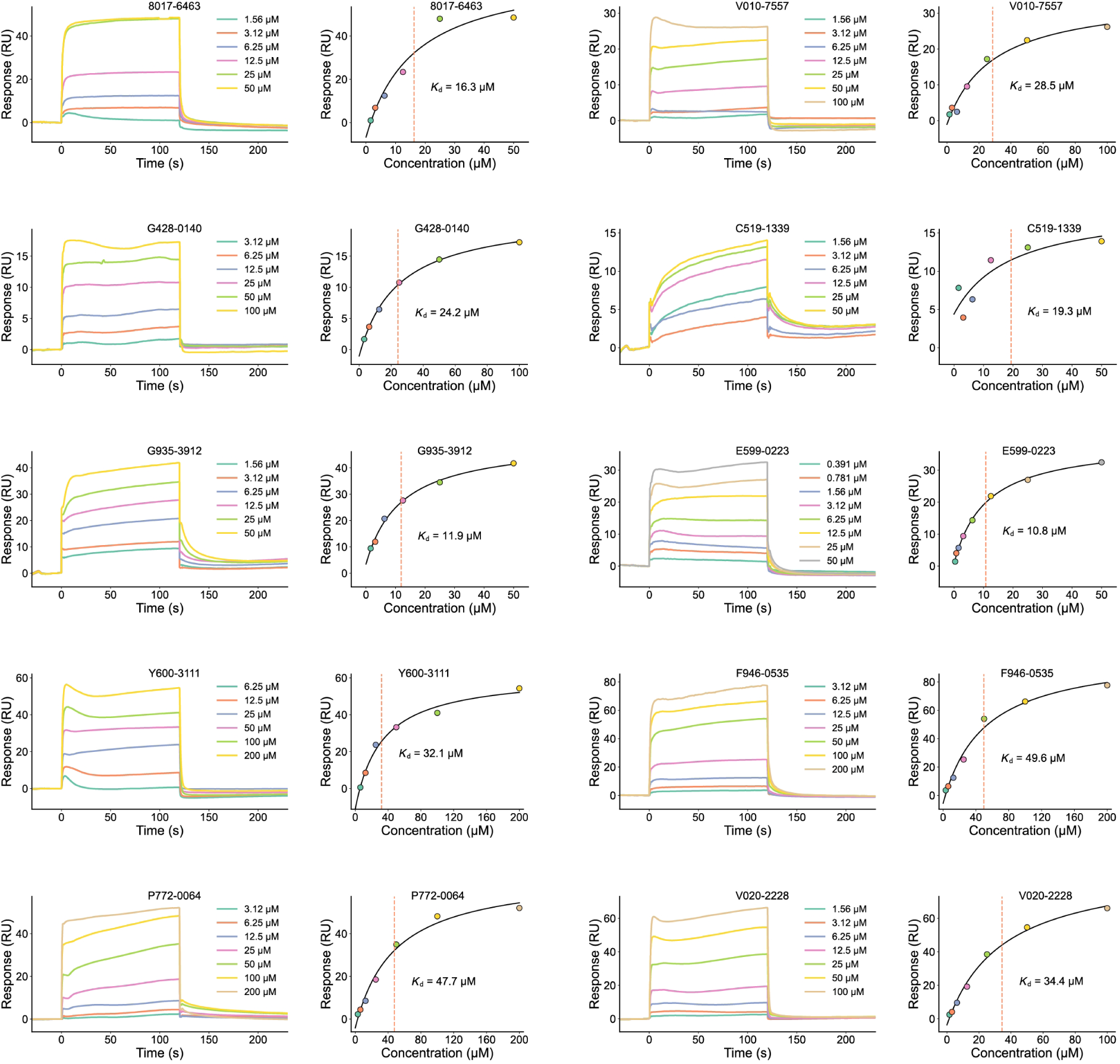
Sensorgrams and steady-state binding curves of the multi-cycle SPR assay for all hit compounds.

**Fig. S12.**
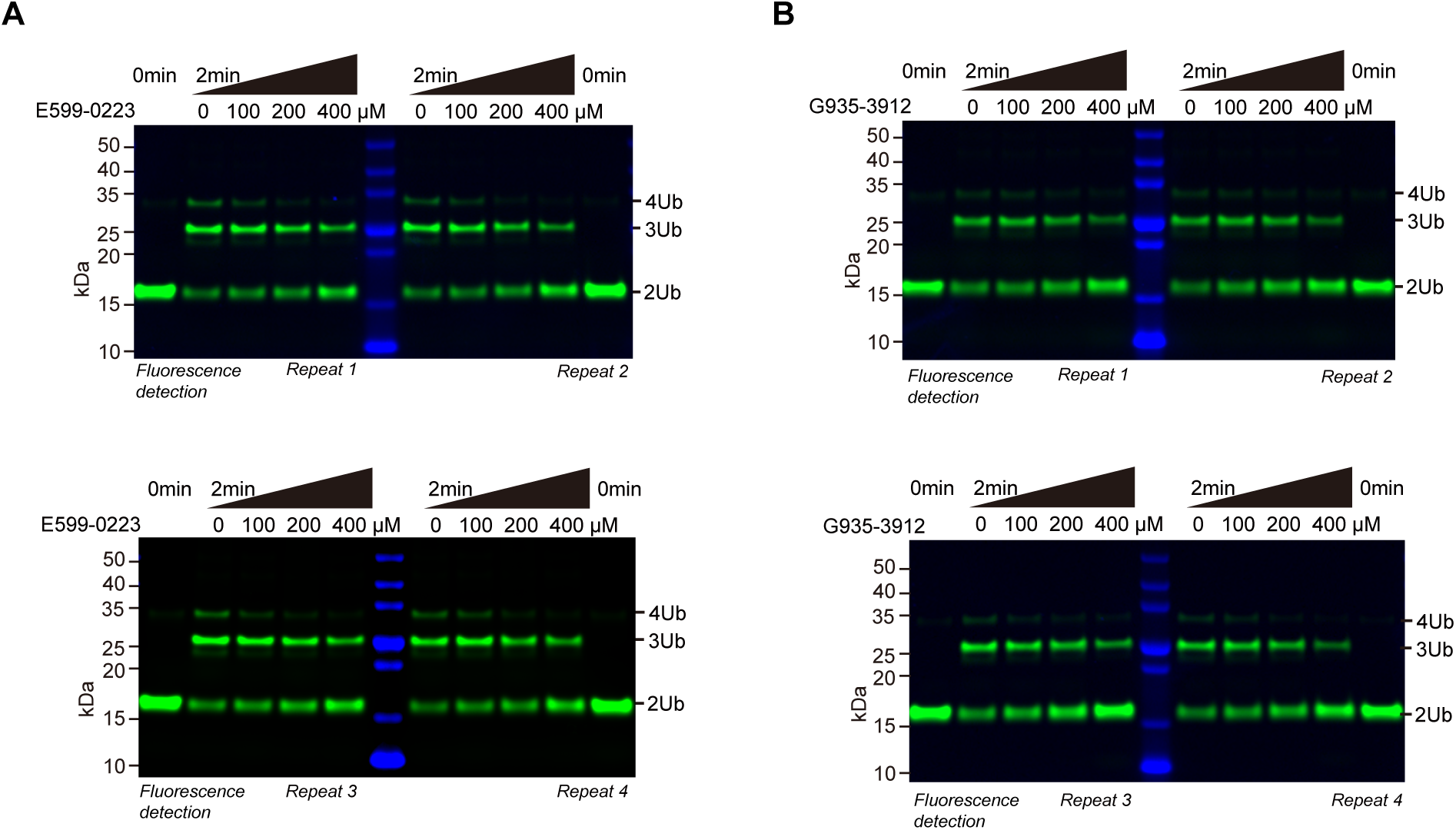
Measuring inhibitory effects of hit compounds to TRIP12 via fluorescent ubiquitination assay. Gel images are representative of independent biological replicates (*n* = 4 for all panels).(A) TRIP12-dependent *in vitro* ubiquitination on fluorescent K48-linked diUb with lysine to arginine mutation at the distal LYS29 site and keeping the proximal LYS29 unchanged (named K48diUb^prox-K29^) with E599-0223. (B) TRIP12-dependent *in vitro* ubiquitination on K48diUb^prox-K29^ with G935-3912.

**Fig. S13.**
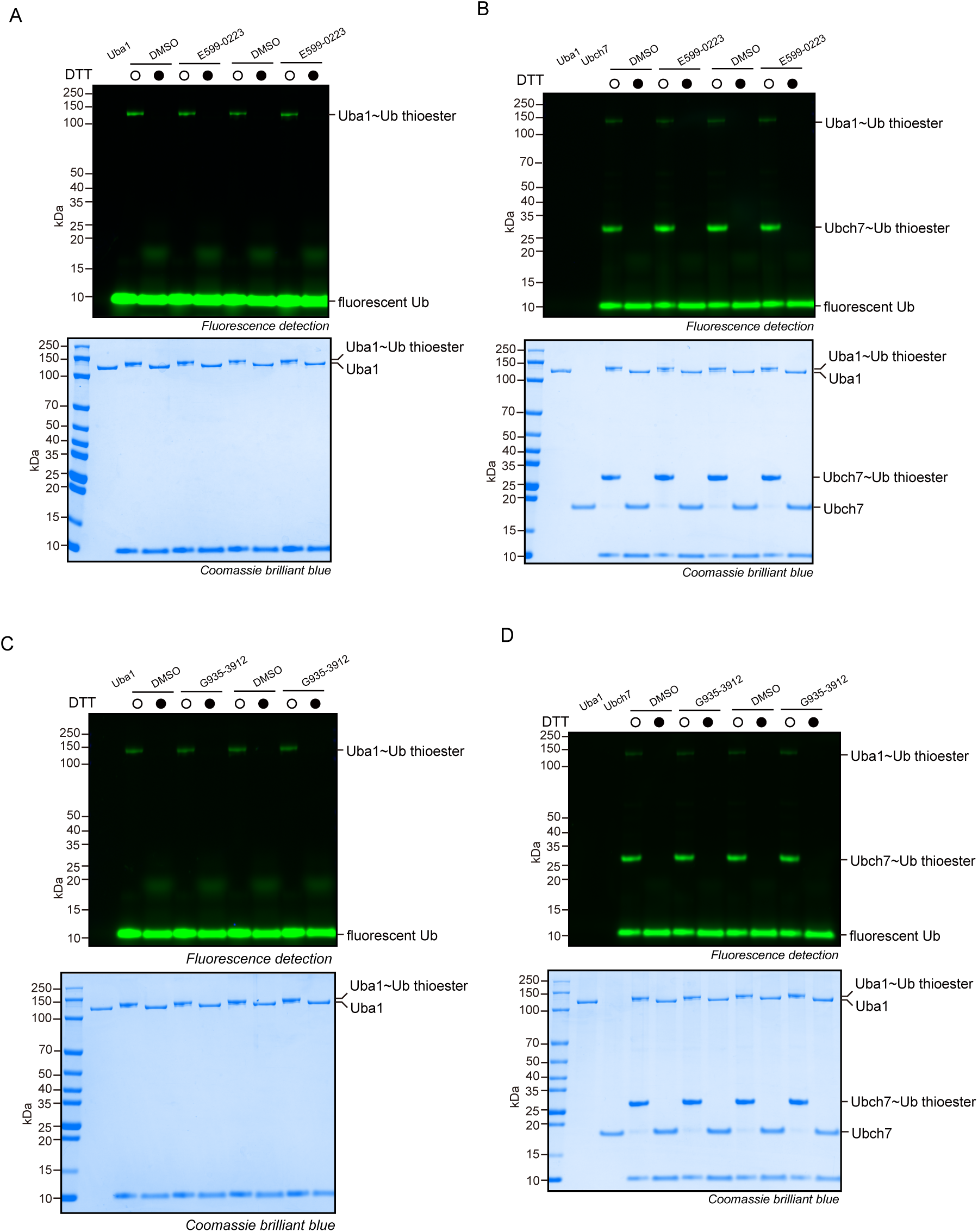
E599-0223 and G935-3912 do not inhibit E1 and E2 enzymes. White circles indicate reactions terminated by SDS, while dark circles indicate reactions terminated by SDS and DTT, which will break thioester bonds. (A) *In vitro* E1∼Ub thioester assay on fluorescent Ub with E599-0223. (B) *In vitro* E2∼Ub thioester assay on fluorescent Ub with E599-0223. (C) *In vitro* E1∼Ub thioester assay on fluorescent Ub with G935-3912. (D) *In vitro* E2∼Ub thioester assay on fluorescent Ub with G935-3912. Gel images are representative of independent biological replicates (*n* = 2 for all panels).

**Table S1.** Druggability prediction results for pocket pretrining, using the RMSE metric.

|  |  | Fpocket ↓ | Druggability ↓ | Total SASA ↓ | Hydrophobicity ↓ |
| --- | --- | --- | --- | --- | --- |
| Finetuning | Uni-Mol | 0.1140 | 0.1001 | 20.73 | 1.285 |
|  | ProFSA | **0.1077** | **0.0934** | **20.01** | **1.275** |
| Zero-shot | Uni-Mol | 0.1419 | 0.1246 | 49.00 | 17.03 |
|  | ProFSA | **0.1228** | **0.1106** | **30.50** | **13.07** |

**Table S2.** Pocket matching results for pocket pretraining, using the AUC metric.

|  | Methods | Kahraman(w/o PO<sub>4</sub>) ↑ | TOUGH-M1 ↑ |
| --- | --- | --- | --- |
| Traditional | SiteEngine | 0.64 | 0.73 |
|  | IsoMIF | 0.75 | - |
| Zero-shot | Uni-Mol | 0.66 | 0.76 |
|  | ProFSA | **0.80** | **0.82** |
| Finetuning | DeeplyTough | 0.67 | 0.91 |
|  | ProFSA | **0.85** | **0.94** |

**Table S3.** Results on LBA prediction task for pocket pertaining, using pearson and spearman correlation

| Method |  | Sequence Identity 30% |  |  | Sequence Identity 60% |  |  |
| --- | --- | --- | --- | --- | --- | --- | --- |
|  |  | RMSE ↓ | Pearson ↑ | Spearman ↑ | RMSE ↓ | Pearson ↑ | Spearman ↑ |
| Sequence Based | DeepDTA | 1.866 | 0.472 | 0.471 | 1.762 | 0.666 | 0.663 |
|  | B&B | 1.985 | 0.165 | 0.152 | 1.891 | 0.249 | 0.275 |
|  | TAPE | 1.890 | 0.338 | 0.286 | 1.633 | 0.568 | 0.571 |
|  | ProtTrans | 1.544 | 0.438 | 0.434 | 1.641 | 0.595 | 0.588 |
| Structure Based | HoloProt | 1.464 | 0.509 | 0.500 | 1.365 | 0.749 | 0.742 |
|  | ATOM3D-3DCNN | 1.416 | 0.550 | 0.553 | 1.621 | 0.608 | 0.615 |
|  | ATOM3D-GNN | 1.601 | 0.545 | 0.533 | 1.408 | 0.743 | 0.743 |
|  | ProNet | 1.463 | 0.551 | 0.551 | <u>1.343</u> | **0.765** | <u>0.761</u> |
| Pretraining Based | GeoSSL | 1.451 | <u>0.577</u> | <u>0.572</u> | - | - | - |
|  | EGNN-PLM | <u>1.403</u> | 0.565 | 0.544 | 1.559 | 0.644 | 0.646 |
|  | Uni-Mol | 1.520 | 0.558 | 0.540 | 1.619 | 0.645 | 0.653 |
|  | ProFSA | **1.377** | **0.628** | **0.620** | **1.334** | <u>0.764</u> | **0.762** |

**Table S4.** Benchmark the performance of DrugCLIP on the DUD-E dataset.

| **Method** | **AUC ↑** | **BEDROC ↑** | **EF1% ↑** |
| --- | --- | --- | --- |
| Vina [62] | 71.60 | – | 7.32 |
| Glide-SP [62] | 76.70 | 40.70 | 16.18 |
| NNScore [63] | 68.30 | 12.20 | 4.02 |
| RF-Score [63] | 65.21 | 12.41 | 4.52 |
| Pafnucy [63] | 63.11 | 16.50 | 3.86 |
| OnionNet [63] | 59.71 | 8.62 | 2.84 |
| PLANET [62] | 71.60 | – | 8.83 |
| GNINA [64] | 76.70 | – | 20.90 |
| DrugCLIP | 77.42 | 39.86 | 24.61 |

**Table S5.** Benchmark the performance of DrugCLIP on the LIT-PCBA dataset.

| **Method** | **AUC ↑** | **BEDROC ↑** | **EF1% ↑** |
| --- | --- | --- | --- |
| Surflex [65] | 51.47 | – | 2.50 |
| Vina [66] | 56.93 | 3.70 | 1.71 |
| Glide-SP [62] | 53.57 | 4.00 | 3.41 |
| NNScore [66] | 55.70 | 2.50 | 1.70 |
| RF-Score [64] | 57.10 | – | 1.67 |
| Pafnucy [67] | – | – | 5.32 |
| PLANET [62] | 55.58 | – | 3.28 |
| GNINA [64] | 61.00 | 5.40 | 4.61 |
| BigBind [68] | 59.07 | – | 3.55 |
| DrugCLIP | 59.54 | 7.29 | 5.36 |

**Table S6.** DUD-E benchmark results with removal of similar molecules from the training set based on ECFP4 similarities and scaffolds.

| Method | AUC ↑ | BEDROC ↑ | EF1% ↑ |
| --- | --- | --- | --- |
| ECFP4 Sim 0.9 | 77.60 | 39.48 | 24.08 |
| ECFP4 Sim 0.6 | 79.02 | 40.82 | 25.27 |
| ECFP4 Sim 0.3 | 77.61 | 31.92 | 19.10 |
| Scaffold | 78.10 | 33.25 | 19.97 |

**Table S7.** DUD-E benchmark results with removal of homologous targets from the training set based on protein sequence similarities and protein families.

| **Method** | **AUC ↑** | **BEDROC ↑** | **EF1% ↑** |
| --- | --- | --- | --- |
| 90% Identity | 77.31 | 39.86 | 24.61 |
| 60% Identity | 75.50 | 32.75 | 19.57 |
| 30% Identity | 73.93 | 29.71 | 17.91 |
| 0% Identity | 69.79 | 16.37 | 9.18 |

**Table S8.** The biochemical and cellular parameters of initially screened positive compounds.

| Compound number | $K_i$ (nM) | β-arr2 NanoBiT | |
| --- | --- | --- | --- |
| | | EC<sub>50</sub> (nM) | $E_{max}$ (%) |
| L589-1477 | 3201.5 | 961.5 | 24.9 |
| F344-0441 | 68.4 | 65.0 | 23.4 |
| 8525-0266 | - | - | - |
| E958-2025 | 138.5 | 163.8 | 14.6 |
| F343-0414 | - | - | - |
| F670-0198 | 1224.2 | 771.2 | 23.0 |
| V006-3328 | 3510.6 | 599.3 | 23.4 |
| V008-4481 | 21.0 | 60.3 | 35.8 |

**Table S9.** Cryo-EM data collection, refinement and validation statistics.

| Category | Y510-9709 | 0086-0043 |
| --- | --- | --- |
| **Data collection and processing** |  |  |
| Magnification | 64,000 | 64,000 |
| Voltage (kV) | 300 | 300 |
| Electron exposure (e<sup>-</sup>/Å<sup>2</sup>) | 50 | 50 |
| Defocus range (μm) | -1.5 to -2.0 | -1.5 to -2.0 |
| Pixel size (Å) | 1.0825 | 1.0825 |
| Symmetry imposed | C2 | C2 |
| Raw movies | 2,918 | 2,687 |
| Particle number | 507 k | 506 k |
| Map resolution (Å) | 2.98 | 2.87 |
| FSC threshold | 0.143 | 0.143 |
| Map resolution range (Å) | 40–2.8 | 40–2.7 |
| **Refinement** |  |  |
| Protein residues | 548 | 548 |
| Ligand | Y510-9709:1 Cl- | 0086-0043:1 Cl- |
| **B factors (Å<sup>2</sup>)** |  |  |
| Protein | 25.76 | 50.53 |
| Ligand | 32.09 | 38.45 |
| Water | 30.28 | 48.95 |
| **R.m.s. deviations** |  |  |
| Bond lengths (Å) | 0.004 | 0.003 |
| Bond angles (°) | 0.666 | 0.631 |
| **Validation** |  |  |
| MolProbity score | 1.64 | 1.41 |
| Clashscore | 6.27 | 5.37 |
| **Ramachandran plot** |  |  |
| Favored (%) | 96.32 | 97.24 |
| Allowed (%) | 3.68 | 2.76 |
| Disallowed (%) | 0.00 | 0.00 |
| **PDB code** | 9JEL | 9JF3 |
| **EMDB code** | EMD-61420 | EMD-61426 |

**Table S10.** The virtual screening performance of DrugCLIP on the DUD-E subset using different pockets on 27 DUD-E targets.

| **Method** | **AUC ↑** | **BEDROC ↑** | **EF1% ↑** |
| --- | --- | --- | --- |
| *holo* - Exp pocket | 81.64 | 46.73 | 29.31 |
| *holo* - fpocket | 78.29 | 39.56 | 23.89 |
| *holo* - fpocket + GenPack | 80.58 | 46.57 | 28.48 |
| AF2 - Exp pocket | 78.56 | 42.27 | 25.88 |
| AF2 - fpocket | 74.47 | 32.11 | 18.96 |
| AF2 - fpocket + GenPack | 79.66 | 39.97 | 24.14 |
| *apo* - Exp pocket | 79.44 | 41.92 | 26.09 |
| *apo* - fpocket | 69.12 | 20.59 | 11.56 |
| *apo* - fpocket + GenPack | 75.59 | 34.16 | 20.43 |

**Table S11.** The virtual screening performance of DrugCLIP on all DUD-E targets with AF2 predictions using different pockets on 96 DUD-E targets.

| **Method** | **AUC ↑** | **BEDROC ↑** | **EF1% ↑** |
| --- | --- | --- | --- |
| *holo* - Exp pocket | 77.31 | 38.88 | 23.97 |
| *holo* - fpocket | 53.72 | 5.87 | 3.19 |
| *holo* - fpocket + GenPack | 75.38 | 34.49 | 20.52 |
| AF2 - Exp pocket | 79.24 | 39.75 | 24.14 |
| AF2 - fpocket | 69.85 | 22.93 | 13.21 |
| AF2 - fpocket + GenPack | 76.28 | 29.43 | 17.02 |

**Table S12.**
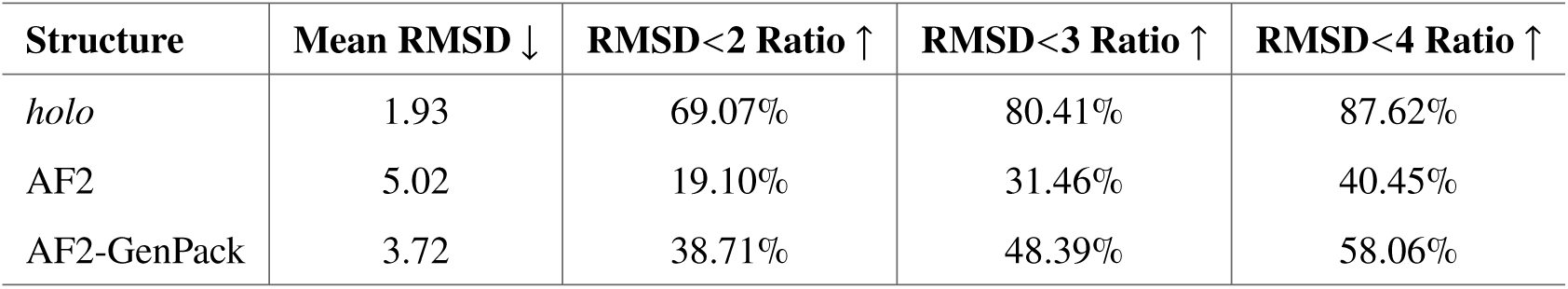
Comparison of mean RMSD and success ratios at different RMSD cutoffs for *holo*, AF2, and AF2-GenPack structures on 96 DUD-E targets.

**Table S13.** Comparison of mean RMSD and success ratios at different RMSD cutoffs for *holo*, AF2, AF2-GenPack, *apo*, and *apo*-GenPack structures on 27 DUD-E targets.

| Structure | Mean RMSD ↓ | RMSD<2 Ratio ↑ | RMSD<3 Ratio ↑ | RMSD<4 Ratio ↑ |
| --- | --- | --- | --- | --- |
| *holo* | 2.57 | 66.67% | 70.37% | 70.37% |
| AF2 | 5.90 | 7.69% | 23.08% | 34.62% |
| AF2-GenPack | 4.41 | 14.81% | 40.74% | 54.15% |
| *apo* | 5.54 | 22.22% | 29.63% | 29.63% |
| *apo*-GenPack | 4.48 | 25.93% | 37.04% | 51.85% |

**Table S14.**
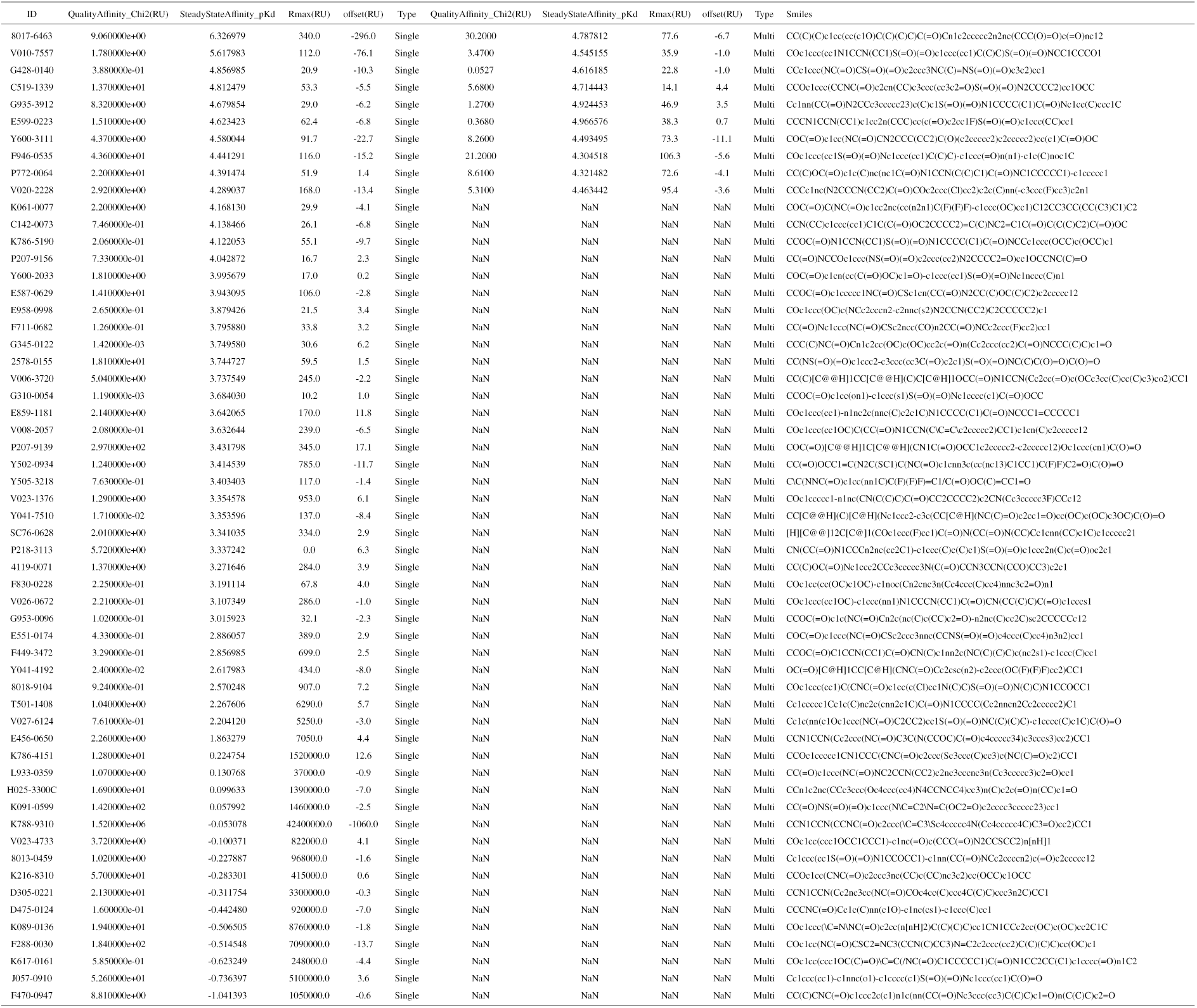
The SPR results of TRIP12 for all wet-lab tested molecules

**Table S15.**
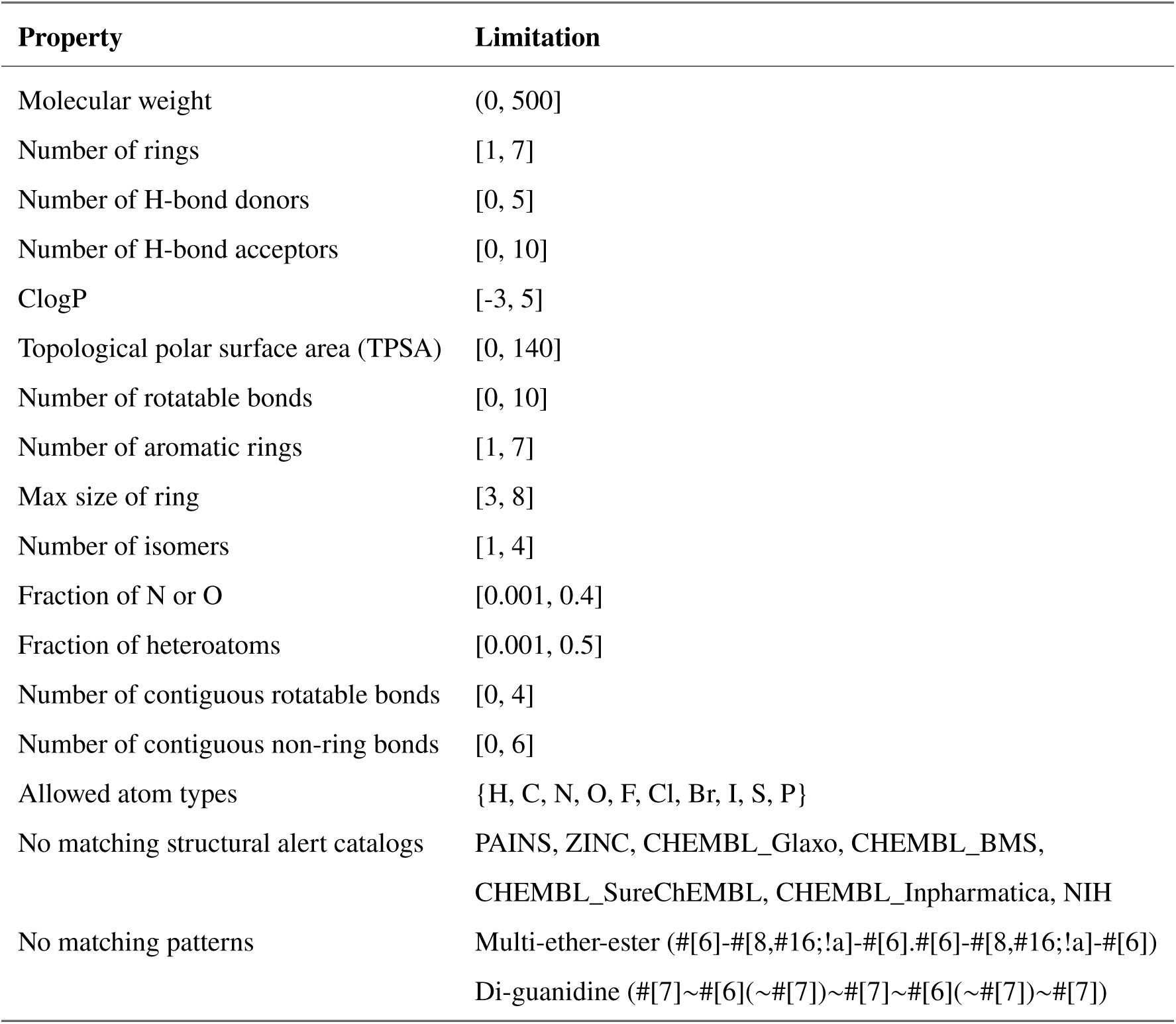
Molecular database filter rules. These rules were concluded based on druglike-ness rules, public structural alerts, and the world (drug) subset of ZINC quantile numbers. Additional constraints on flexibility-related properties were imposed to prevent a sharp increase in the computational cost of molecular docking.

## Notes

https://drug-the-whole-genome.yanyanlan.com/

### Competing Interest Statement

The authors have declared no competing interest.

### Summary of Updates

First, we added new experimental validation on TRIP12, a challenging and previously uncharacterized target with no known ligands or structural data. In Fig. 3 of this updated version, we present a hit rate of >15% by DrugCLIP screening solely on AlphaFold2 predicted structures of the HETC domain of TRIP12. SPR (surface plasmon resonance) results confirmed a direct physical interaction between our hits and TRIP12, while fluorescent ubiquitination assays proved the functional inhibitory effect. These results address concerns regarding DrugCLIP ability to generalize beyond well-studied proteins and provide strong evidence for its applicability to the underexplored regions of the proteome. Second, we conducted additional analyses to explore why GenPack improves downstream virtual screening performance. Our results show that enhanced binding site localization is the main factor driving improved screening accuracy, rather than side-chain RMSD or overall structural refinement. These findings clarify the mechanistic underpinnings of GenPack and address concerns regarding the relationship between structural accuracy and screening performance. Finally, we expanded the Methods section to provide greater clarity and transparency. We now include detailed descriptions of the training set split for the GenPack model, the compound selection and filtering process for DrugCLIP-based screening, and the data construction and augmentation strategy for the ProFSA and DrugCLIP pipeline. These updates provide more technical details and help ensure reproducibility.

